# The Dystonia Gene THAP1 Controls DNA Double Strand Break Repair Choice

**DOI:** 10.1101/2020.07.19.210773

**Authors:** Kenta Shinoda, Dali Zong, Elsa Callen, Wei Wu, Lavinia C. Dumitrache, Frida Belinky, Nancy Wong, Momoko Ishikawa, Andre Stanlie, Michelle Ehrlich, Peter J. McKinnon, Andre Nussenzweig

**Affiliations:** Laboratory of Genome Integrity, National Cancer Institute, NIH, Bethesda, MD, USA; St. Jude Translational Neuroscience, Center for Pediatric Neurological Disease Research, Depts. Genetics & Cell Mol. Biology, St. Jude Children’s Research Hospital, Memphis, TN, USA; Department of Neurology, Icahn School of Medicine at Mount Sinai, New York, NY, USA

## Abstract

The Shieldin complex, consisting of SHLD1, SHLD2, SHLD3 and REV7, shields DNA double strand breaks (DSBs) from nucleolytic resection. The end-protecting activity of Shieldin promotes productive non-homologous end joining (NHEJ) in G1 but can threaten genome integrity during S-phase by blocking homologous recombination (HR). Curiously, the penultimate Shieldin component, SHLD1 is one of the least abundant mammalian proteins. Here, we report that the transcription factors THAP1, YY1 and HCF1 bind directly to the *SHLD1* promoter, where they cooperatively maintain the low basal expression of *SHLD1*. Functionally, this transcriptional network ensures that SHLD1 protein levels are kept in check to enable a proper balance between end protection and end resection during physiological DSB repair. In the context of BRCA1 deficiency, loss of THAP1 dependent SHLD1 expression confers cross resistance to PARP inhibitor and cisplatin, and shorter progression free survival in ovarian cancer patients. In contrast, loss of THAP1 in BRCA2 deficient cells increases genome instability and correlates with improved responses to chemotherapy. Pathogenic THAP1 mutations are causatively linked to the adult-onset torsion dystonia type 6 (DYT6) movement disorder, but the critical disease targets are unknown. We further demonstrate that murine models of *Thap1*-associated dystonia show reduced *Shld1* expression concomitant with elevated levels of unresolved DNA damage in the brain. In summary, our study provides the first example of a transcriptional network that directly controls DSB repair choice and reveals a previously unsuspected link between DNA damage and dystonia.

---

Directjoining of double strand breaks (DSBs) by non-homologous end joining (NHEJ) provides eukaryotes with a fast and versatile mechanism for repairing genomic damage ^1^. Although not as precise as homologous recombination (HR), NHEJ is essential for genome maintenance in non-dividing cells (such as post-mitotic neurons), in which sister chromatid templates for high fidelity HR are unavailable. In addition, NHEJ supports physiological DNA rearrangement processes where genetic diversity is desirable, such as V(D)J recombination and immunogolobulin class switching (CSR) in developing lymphocytes ^2^. Accordingly, inactivation of NHEJ factors have been linked to neurodegeneration and immunodeficiency in model organisms as well as in human patients ^3–5^.

Once cells enter S-phase to commence DNA replication, HR gradually becomes an option as sister chromatids become available ^6^. Extensive 5’-nucleolytic resection of DSBs is required to license HR, and is thought to simultaneously preclude NHEJ. The 3’-single-stranded (ss) overhang generated by resection is initially stabilized by RPA. The PALB2/BRCA2 mediator complex subsequently displaces RPA from the ssDNA and assembles polymers of the recombinase RAD51, which invade the intact sister chromatid to promote templated DNA synthesis ^7^. HR is uniquely suitable for healing replication-associated DSBs, as most of them are one-ended. By contrast, NHEJ is either unable to repair such lesions or would give rise to translocations when disparate one-ended DSBs are inappropriately joined together. Consistently, deficiencies in HR not only facilitate tumorigenesis ^8^, but also confer hypersensitivity to chemotherapeutic agents such as PARP inhibitors (PARPi) and cisplatin ^9^.

53BP1 has emerged as a master coordinator of anti-HR activities in mammalian cells ^10, 11^. 53BP1 recruits a number of effector molecules, including PTIP ^12^, the Shieldin complex (consisting of SHLD1, SHLD2, SHLD3 and REV7) ^13–22^, the CST complex ^19, 23^ and DYNLL1 ^24^, to block DSB end resection by multiple nucleases. In addition, the 53BP1-RIF1-Shieldin subpathway is capable of directly impeding PALB2/BRCA2- mediated RAD51 loading on ssDNA ^25^. These barriers are normally overcome by the tumor suppressor BRCA1, which counteracts 53BP1 during S-phase by stimulating both nucleolytic resection and RAD51 assembly through still poorly defined mechanisms ^8, 26^. In addition to BRCA1, the AAA+ family ATPase TRIP13 promotes HR by catalyzing an inactivating conformational change in the REV7 component of the Shieldin complex ^27^.

Much of the HR defects observed in BRCA1-deficient cells can be attributed to excessive 53BP1-mediated end-protection, which in turn promotes the toxic joining of replication-associated DSBs ^25, 28–30^. Notably, depletion of any Shieldin complex component restores HR in BRCA1-deficient cells ^14, 16–21^, and reduced expression of Shieldin is associated with the acquisition of PARPi resistance in PDX models of BRCA1- deficient breast cancer ^14^.

These observations suggest that the level of Shieldin activity is a major determinant of HR competency and chemosensitivity in mammalian cells. Remarkably, however, Shieldin components are undetectable by mass spectrometry based proteome analysis ^18^ and are amongst the lowest expressed genes in the genome ^31^. Here, using a combination of CRISPR screens and biochemical assays, we addressed how changes in the levels of Shieldin affect DSB repair and genome stability. Our work reveals that the basal expression of SHLD1 is maintained by direct binding of a network of transcription factors consisting of THAP1, HCF1 and YY1 to its promoter. Mutations in *THAP1* cause an inherited form of dystonia (DYT6), a neurodegenerative disorder that is characterized by sustained involuntary muscle contractions ^32–34^, although critical disease target(s) of THAP1 are unknown. We find that inactivation of THAP1 abolishes *SHLD1* expression, leading to HR restoration and chemoresistance in BRCA1-deficient cell lines and BRCA1- mutated human ovarian cancers. Conversely, overexpression of either SHLD1 or THAP1 compromises the repair of replication-associated DSBs in BRCA1-proficient cells. Moreover, THAP1 is essential for CSR and helps maintain genome integrity in the developing brain. Together, these results help explain why Shieldin expression must be kept at low levels, and reveal a hitherto unknown physiological function of THAP1 in maintaining genome stability in all cell types.

## Results

### Identification of THAP1 as a modifier of chemosensitivity in mouse models of BRCA1 deficiency

The Shieldin complex is recruited to DSBs by 53BP1 and RIF1 where it blocks HR by inhibiting nucleolytic end resection ^10, 11^ as well as suppressing the loading of RAD51 onto ssDNA post-resection ^25, 35^. These barriers to HR are normally overcome by BRCA1, but become insurmountable in BRCA1-deficient cells and cause PARP inhibitor hypersensitivity. Here, we undertook whole-genome CRISPR-Cas9 screens to search for genes whose mutation confer PARPi resistance in BRCA1-deficient cells. Human BRCA1-deficient tumors, particularly those that harbor pathogenic exon 11 mutations, often express a hypomorphic BRCA1 protein (BRCA1-Δ11q) ^36^. In order to model acquired PARPi resistance in a clinically relevant manner, we elected to use the PARPi-hypersensitive *Brca1*^Δ*11*^ and *Brca1*^Δ*11*^*Trp53bp1^S25A^* murine models, in which a truncated BRCA1 protein (BRCA1-Δ11) highly similar to human BRCA1-Δ11q is expressed ^25, 29, 36^ (Fig. 1a). Moreover, cells derived from *Brca1*^Δ*11*^ and *Brca1*^Δ*11*^*Trp53bp1^S25A^* mice exhibit marked HR defects due to excessive Shieldin-mediated end protection pre- and post-resection, respectively ^25^, thereby representing murine model systems to search for putative regulators of Shieldin function.

**Fig. 1.**
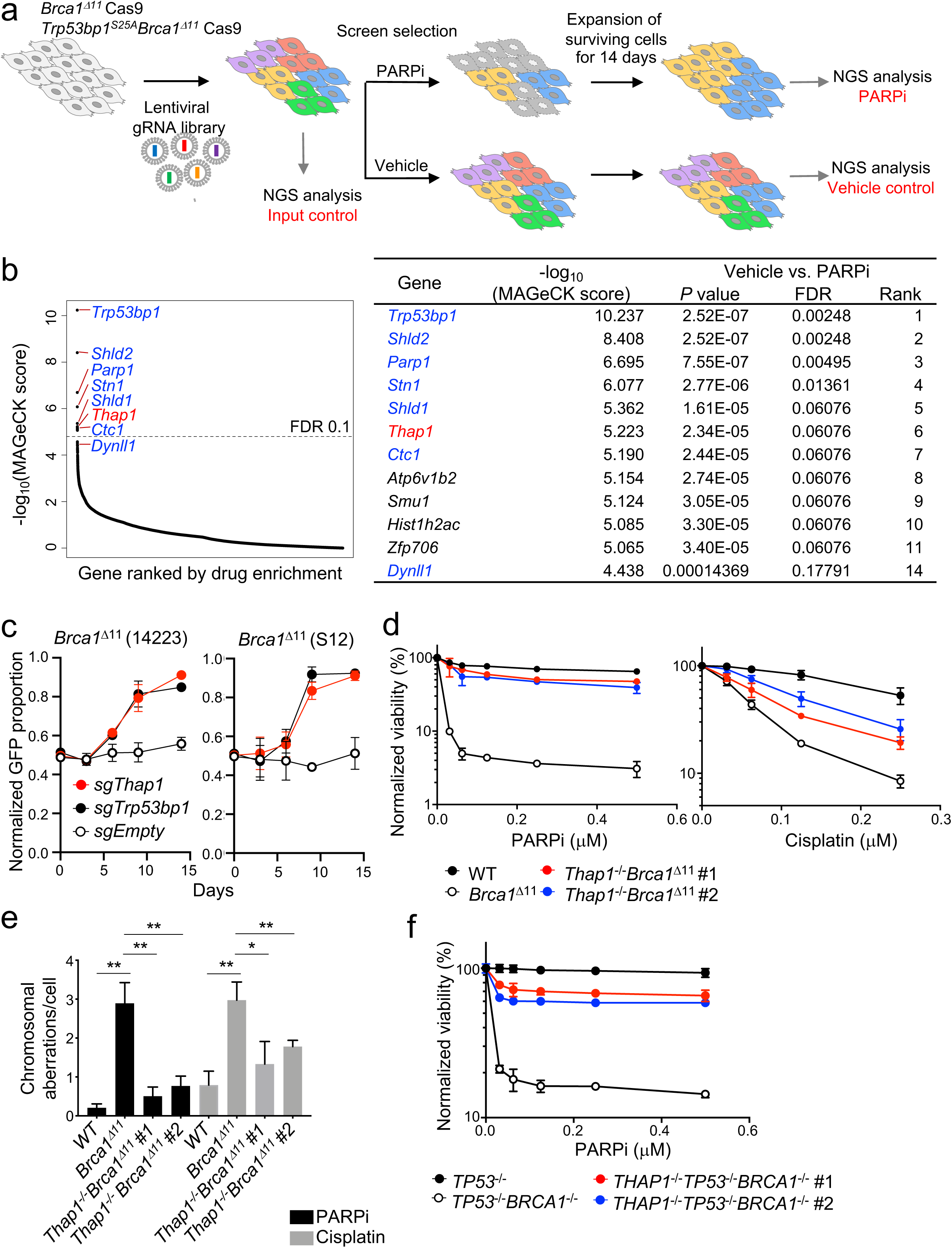
**Identification of THAP1 as a modifier of chemosensitivity in BRCA1-deficient cells.** (**a**) Schematic of CRISPR-based screen for genes whose deletion confer resistance to the clinical PARP inhibitor (PARPi) olaparib in *BRCA1*-mutant MEFs (*Brca1*^Δ*11*^and *Brca1*^Δ*11*^*Trp53bp1^S25A^*). (**b**) Analysis of guide RNA enrichment in PARPi survivors by MAGeCK. A false discovery rate (FDR) of 0.1 is indicated by dotted line. The combined top hits for *Brca1*^Δ*11*^ (n=2) and *Brca1*^Δ*11*^*Trp53bp1^S25A^* (n=1) screens are tabulated on the right. summary of n = 3 biological replicates. Previously identified resistance genes are shown in blue and *Thap1* is shown in red. (**c**) Multicolor Competition Assay (MCA): *Cas9^+^Brca1*^Δ*11*^ MEFs (14223 and S12) transduced with a specific guide RNA targeting *Thap1*, *Trp53bp1* or an empty vector (sgEmpty, all GFP-positive) were co-incubated (1:1 ratio) with *Cas9^+^Brca1*^Δ*11*^ MEFs transduced with non-targeting guides (sgLacZ, mCherry-positive). Data represent mean fraction of GFP-positive cells ± s.d., normalized to day 0 (n = 3). Deletion of *Thap1* significantly enhanced the outgrowth of *Brca1*^Δ*11*^ cells following PARPi (100 nM) treatment (p<0.0001). (**d**) Viability of *WT*, *Brca1*^Δ*11*^ (14223) and two individual clones (#1 and #2) of *Thap1^-/-^ Brca1*^Δ*11*^ MEFs (derived from 14223), as measured by CellTiter-Glo seven days after treatment with PARPi (left) and cisplatin (right). (**e**) Genomic instability detected in the metaphase spreads of *WT*, *Brca1*^Δ*11*^ and two individual clones (#1 and #2) of *Thap1^-/-^Brca1*^Δ*11*^ MEFs. Cells were treated for 16 h with 1 µM PARPi or with 0.5 µM cisplatin. At least 50 metaphase spreads were scored per genotype and condition. The experiments were repeated four (PARPi) and three (cisplatin) times, respectively. Statistical significance was determined by Welch’s t-test. (**f**) Viability of *TP53^-/-^*, *TP53^-/-^BRCA1^-/-^* and two individual clones (#1 and #2) of *THAP1^-/-^ TP53^-/-^BRCA1^-/-^* human RPE1 cells, as measured by CellTiter-Glo seven days after PARPi treatment.

*Brca1*^Δ*11*^ and *Brca1*^Δ*11*^*Trp53bp1^S25A^* mouse embryonic fibroblasts (MEFs) expressing CAS9 were transduced with lentiviral libraries of single-guide RNAs (sgRNAs), and the resultant pools of edited cells were exposed to near-lethal doses of the clinically used PARPi olaparib for two weeks (Fig. 1a). The screens were performed twice in *Brca1*^Δ*11*^ MEFs and once in *Brca1*^Δ*11*^*Trp53bp1^S25A^* MEFs. Gene enrichment or depletion was calculated for each cell line using the MAGeCK algorithm ^37^, with positive and negative b-scores indicating positive and negative selection of a given gene deletion event, respectively (Fig. 1b and Table S1). To obtain high-confidence hits, we combined the individual gene-based b-scores for both cell lines. Importantly, in both cell lines we identified multiple genes whose deletion have previously been shown to confer PARPi resistance in human BRCA1-deficient cells, including *Trp53bp1, Shld1*, *Shld2*, *Ctc1*, *Stn1* and *Dynll1* (Fig. 1b and Table S1) ^14–21, 24^, thereby confirming the validity of our approach.

In addition, we identified *Thap1* as a candidate gene whose deletion strongly imparts PARPi resistance in both *Brca1*^Δ*11*^ and *Brca1*^Δ*11*^*Trp53bp1^S25A^* MEFs (Fig. 1b). *Thap1* encodes the murine homolog of Thanatos-associated protein 1 (THAP1), a transcription factor that is causatively linked to adult-onset torsion dystonia type 6 (DYT6) in humans ^32-34^. To verify the results of the primary screens, we conducted competitive growth assays using sgRNAs targeting either *Thap1* or *Trp53bp1* and found that each led to the outgrowth of two independently-derived *Brca1*^Δ*11*^ MEF cell lines (14223 and S12) treated with PARPi (Fig. 1c and Extended Data Fig. 1a), while no difference in growth was observed without the drug (Extended Data Fig. 1b). In addition, we generated two independent clonal knockouts of *Thap1* in one of the two *Brca1*^Δ*11*^ MEF cell lines (14223) (Extended Data Fig. 1c) and confirmed that deletion of *Thap1* led to cross-resistance to PARPi and cisplatin, as measured by cell viability assays and mitotic chromosome aberrations (Fig. 1d, e). Finally, deletion of *THAP1* in two-independently derived clones of *BRCA1*-null human RPE1 cells ^20^ resulted in marked PARPi resistance (Fig. 1f). In contrast to BRCA1 deficiency, *Thap1* deletion did not affect the PARPi sensitivity of BRCA1-proficient cells as measured by cell viability (Extended Data Fig. 1d, e). These results support THAP1 as a *bona fide* modifier of chemosensitivity in mouse and human cell lines with BRCA1 deficiency.

### Loss of THAP1 confers chemoresistance in BRCA1-deficient human tumors

Analysis of publicly available gene expression databases revealed that *THAP1* is ubiquitously expressed in malignant tissues, with notable increased expression in testis, ovarian and breast cancers (Fig. 2a). To test whether dysregulated *THAP1* has an impact on tumorigenesis, we extracted human tumor expression data from The Cancer Genome Atlas (TCGA) database ^38^. We found that low expression of *THAP1* was significantly correlated with shorter progression-free survival (PFS) of patients with *BRCA1*-mutated serous ovarian carcinoma (Fig. 2b). Although most patients in this cohort have undergone platinum-based chemotherapy ^38^, no correlation with PFS was observed for patients with *BRCA1*-proficient tumors (Extended Data Fig. 2a). Together, our results suggest that *THAP1* loss alleviates PARPi and cisplatin hypersensitivity in BRCA1-deficient cells and tumors, and this effect is not species-, cell lineage- or *BRCA1* mutation-specific.

**Fig. 2.**
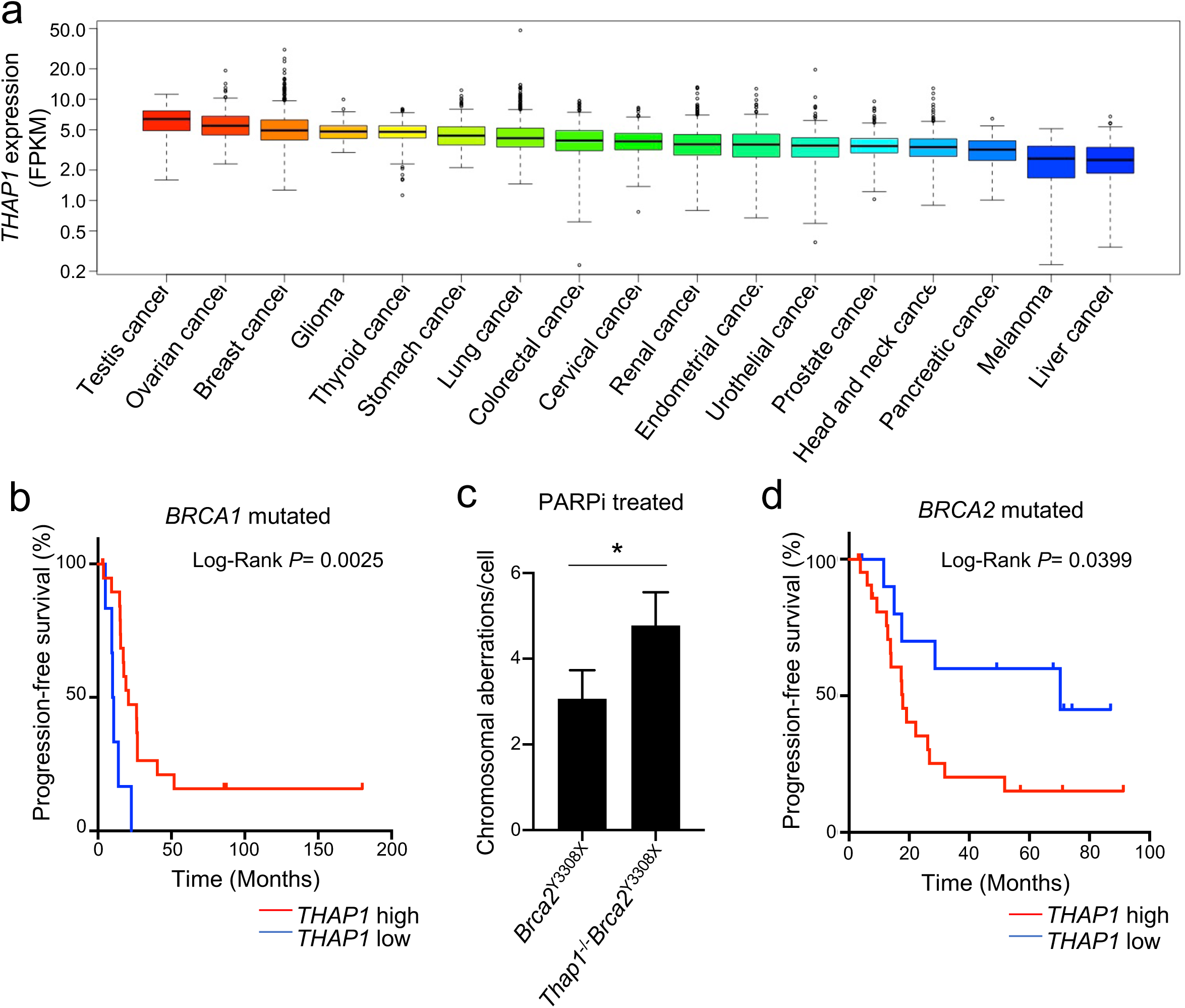
**Loss of THAP1 expression affects the chemosensitivity of BRCA-deficient human tumors.** (**a**) Expression of the *THAP1* gene (FPKM) in 17 different cancer types. Data were obtained from The Cancer Genome Atlas (TCGA) project. (**b**) Progression-free survival (PFS) of BRCA1-mutated ovarian serous adenocarcinoma patients with standard platinum-based regimens ^38^. Patients were defined as having *THAP1* low- or high-expressing tumors on the basis of the quintile of *THAP1* expression (z-scores < −0.67). PFS curves were generated by the Kaplan–Meier method. The difference between the PFS of *THAP1* low- versus *THAP1* high-expressing patients was assessed by the log-rank test (P < 0.01). (**c**) Genomic instability detected in the metaphase spreads of *Brca2^Y3308X^* and *Thap1^-/-^ Brca2^Y3308X^* mouse embryonic ES cells (mESCs) after 16 h of PARPi treatment (0.5 µM). At least 50 cells were scored per genotype and condition. The experiment was repeated three times. Statistical significance was determined by Welch’s t-test. (**d**) PFS of BRCA2-mutated ovarian serous adenocarcinoma patients with standard platinum-based regimens ^38^. Patients were defined as having *THAP1* low- or high-expressing tumors on the basis of the same *THAP1* expression (z-scores < −0.67) used in Fig. 2b. PFS curves were generated by the Kaplan–Meier method. The difference between the PFS of *THAP1* low-versus *THAP1* high-expressing patients was assessed by the log-rank test (P < 0.05).

A recent study of prognostic biomarkers of acute myelogenous leukemia reported that low levels of *BRCA2* and *THAP1* correlated with improved patient survival ^39^. Since BRCA1 and BRCA2 have functionally non-redundant roles during HR, we also sought to determine how *THAP1* expression influences the response of BRCA2-deficient cells to chemotherapy. In contrast to its impact on BRCA1-deficiency (Fig. 1c-f and 2b), we found that deletion of *Thap1* increased genome instability in *Brca2^Y3308X^* mutant mouse embryonic stem cells (mESCs) treated with PARPi (Fig. 2c). Consistent with increased therapeutic vulnerability, clinical data demonstrated that low *THAP1* expression was correlated with longer PFS in patients with *BRCA2*-mutant serous ovarian carcinoma (Fig. 2d). In summary, our results indicate that expression of THAP1 modulates the responses to chemotherapy in opposite ways in BRCA1- and BRCA2-deficient cells (see discussion).

### THAP1 is a direct transcriptional regulator of *Shld1* expression

Mutations in *THAP1* have been identified as the cause of DYT6, which is associated with abnormal muscle contractions without overt neuropathological lesions ^32–34^. THAP1 has been proposed to regulate numerous targets in neurons, but the causative links to dystonic movements remains unclear. While several transcription factors promote DNA repair by acting directly at sites of DNA damage in a transcription-independent manner ^40^, we failed to observe THAP1 translocation to sites of DNA damage induced by g-irradiation (not shown). Given that most of the dystonia-causing mutations in THAP1 disrupt its sequence-specific DNA binding activity ^41, 42^, we therefore hypothesized that THAP1 may regulate the chemosensitivity of BRCA-deficient cells directly through transcriptional mechanisms.

To investigate this possibility, we performed nascent RNA-sequencing (RNA-seq) to systematically probe for genes that are differentially expressed in *Thap1*^-/-^ and *Thap1*^-/-^ *Brca1*^Δ*11*^ MEFs, as compared to wild-type and *Brca1*^Δ11^ parental cells, respectively. In total, we found 452 differentially expressed genes (221 downregulated and 231 upregulated (log2 fold-change >2 and FDR <0.05) between *Thap1*^-/-^ and wild-type cells, while 1,337 genes were differentially expressed (675 downregulated and 662 upregulated) between *Thap1*^-/-^*Brca1*^Δ*11*^ and *Brca1*^Δ*11*^ cells (Fig. 3a). Of these, 98 genes (57 downregulated and 41 upregulated) were common to both *Thap1*^-/-^ and *Thap1*^-/-^*Brca1*^Δ11^ cells (Table S2). To determine which of these genes are direct targets of THAP1, we performed THAP1 ChIP-sequencing (ChIP-seq) in MEFs which identified 2,137 unique THAP1 binding sites. Overall, only six differentially expressed genes in *Thap1*-deficient MEFs were also bound by THAP1 (Fig. 3b). Among these genes we identified *Shld1*, the penultimate component of the Shieldin complex and a known effector of 53BP1. Re-examining our RNA-seq datasets, we confirmed that *Shld1* expression was significantly decreased in *Thap1^-/-^* and *Thap1^-/-^BRCA1*^D^*^11/^*^Δ*11*^ MEFs (Fig. 3c). By contrast, other genes encoding 53BP1 pahway members exhibited only modest expression changes (Extended Data Fig. 3a, b).

**Fig. 3.**
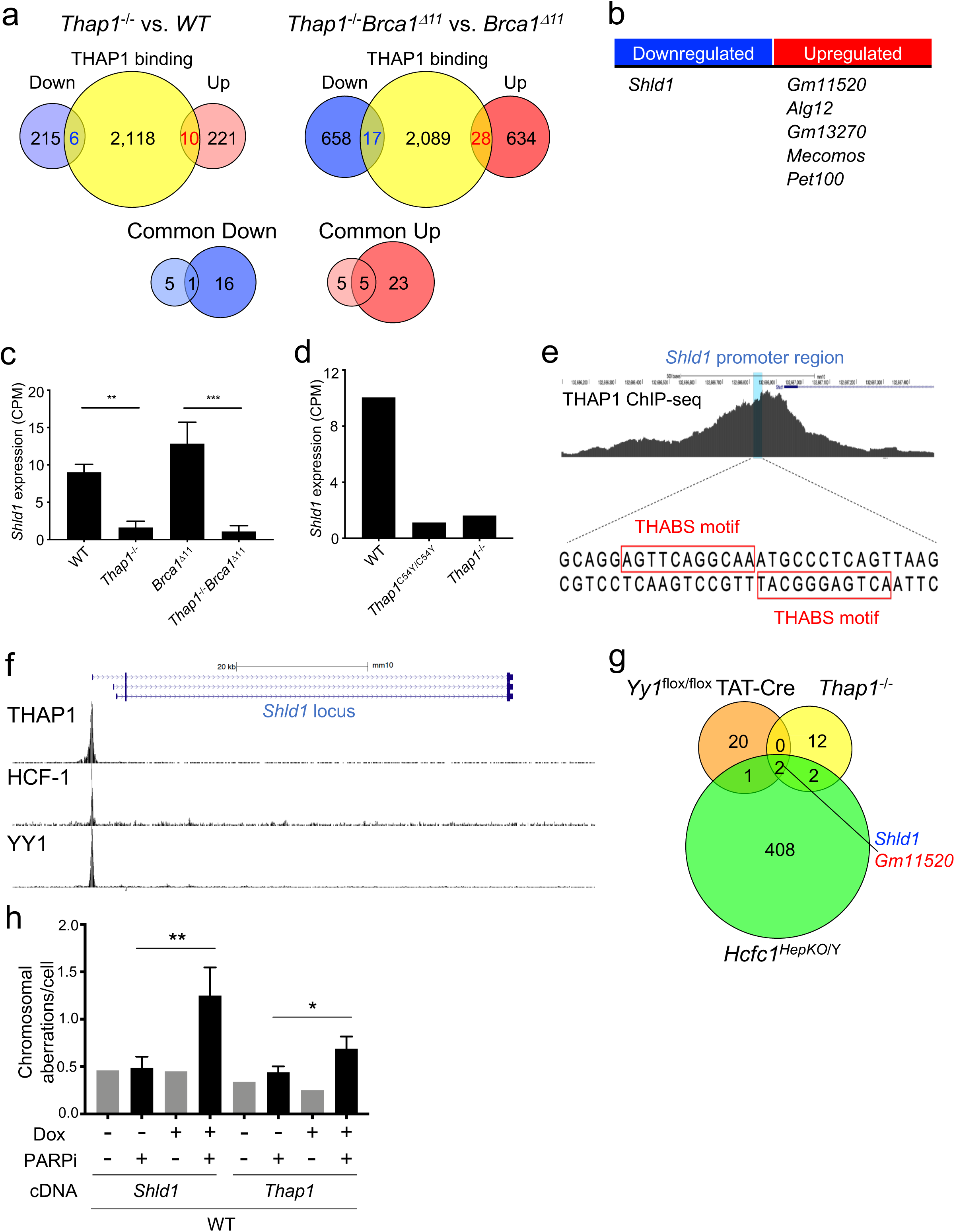
**THAP1 co-regulates *Shld1* expression with HCF1 and YY1.** (**a**) Top: Venn diagram depicting differentially expressed genes (nascent RNA-seq, log2 fold-change >2 and FDR <0.05) in *Thap1^-/-^* versus *WT* and *Thap1^-/-^Brca1*^Δ*11*^ versus *Brca1*^Δ*11*^ MEFs in relation to THAP1-bound genes (ChIP-seq). The number of genes that were shown to be bound by THAP1 and were either downregulated or upregulated in THAP1-deficient MEFs are shown in blue and red, respectively. Bottom: Venn diagram showing the number of differentially expressed THAP1-bound genes that were common to both *Thap1^-/-^* and *Thap1^-/-^Brca1*^Δ*11*^ MEFs. (**b**) List of differentially expressed THAP1-bound genes that were common to both *Thap1^- /-^* and *Thap1^-/-^Brca1*^Δ*11*^ MEFs. (**c**) Levels of *Shld1* gene expression in *WT, Thap1^-/-^, Brca1*^Δ*11*^ and *Thap1^-/-^Brca1*^Δ*11*^ MEFs, as detected by nascent RNA-seq. Data represent mean counts per million reads (CPM) ± s.d. (n = 3, **p<0.01, ***p<0.001). Statistical significance were determined by one-way ANOVA. (**d**) Levels of *Shld1* gene expression in *WT, Thap^C54Y/C54Y^ and Thap1^-/-^* mESCs. Data are from a publicly available RNA-seq dataset (GSE86911). Data represent mean CPM. (**e**) The location of the THAP1 binding sequence (THABS) motif within the *Shld1* promoter. THAP1 ChIP-seq signal at the Shld1 promoter region is shown above. (**f**) Binding of THAP1, HCF1 and YY1 at the *Shld1* gene locus. Data are from publicly available ChIP-seq datasets in ES cells (THAP1, GSE86911; HCF1, GSE36030; YY1, GSE68195). (**g**) Venn diagram depicting differentially expressed genes (log2 fold-change >2 and FDR <0.05) in *Thap1^-/-^* MEFs, *Yy1^flox/flox^* TAT-Cre, and *Hcfc1^HepKO/Y^* cells in relation to THAP1-bound genes and previously published YY1- and HCF1-bound genes. Only two genes bound by all three transcription factors also exhibited differential expression: *Shld1* (downregulated) and *Gm11520* (upregulated). (**h**) Genomic instability detected in the metaphase spreads of *WT* MEFs overexpressing SHLD1 or THAP1 with or without 16 h of PARPi treatment (1 µM). Ectopic SHLD1 and THAP1 expression was induced by doxycycline (Dox) treatment. At least 50 cells were scored per genotype and condition. The experiments were repeated four times for PARPi treated condition and twice for untreated condition. Statistical significance was determined by Welch’s t-test.

Sequence-specific DNA binding of THAP1 is mediated by the THAP-type zinc finger located within its THAP domain and is disrupted by the C54Y point mutation, a known DYT6 causative mutation ^43, 44^. To determine whether THAP1 DNA binding is required for *Shld1* expression, we re-analyzed a RNA-seq dataset derived from *Thap1^C54Y/C54Y^* mESCs ^45^. We found that *Shld1* expression was decreased in *Thap1^C54Y/C54Y^* cells at a level comparable to that found in *THAP1*^-/-^ mESCs (Fig. 3d and Extended Data Fig. 3c). Thus, sequence-specific DNA binding is required for THAP1 regulation of SHLD1.

### A THAP1-HCF1-YY1 co-regulatory module promotes *Shld1* expression

The *Shld1* promoter contains a consensus THAP1 binding sequence (THABS) motif where a strong THAP1 ChIP-seq signal was detected (Fig. 3e). Recent studies have shown that THAP1 frequently co-occupy its target genes with the transcriptional co-regulators HCF1 and YY1 ^46–48^, which are recruited to the promoters by THAP1 ^47, 49^. Analyses of ChIP-seq datasets revealed that HCF1 and YY1 indeed co-occupy the *Shld1* promoter with THAP1 (Fig. 3f). Moreover, we found that inducible deletion of *Hcfc1* in hepatocytes (*Hcfc1^HepKO/Y^*) led to downregulation of *Shld1* expression (Extended Data Fig. 3d) ^50^. Similarly, conditional knockout of *Yy1* in mouse B cells (*Yy1^flox/flox^*TAT-Cre) significantly decreased *Shld1* expression (Extended Data Fig. 3e) ^51^. Strikingly, when we overlapped genes whose expression were differentially regulated by HCF1, YY1 and THAP1 and that were also direct targets of these transcription factors, only two genes *Shld1* and *Gm11520* (a putative noncoding RNA of unknown function) showed significant enrichment (Fig. 3g and Extended Data Fig. 3f, g). Thus, although the vast majority of genes appear to be independently regulated by THAP1, HCF1 and YY1, these cofactors cooperatively bind to the *Shld1* promoter to maintain its expression.

Given that Shld1 has been shown to be amongst the lowest expressed genes ^18, 52^, we next investigated whether elevating SHLD1 protein levels cause cellular toxicity. To this end, we overexpressed exogenous SHLD1 or THAP1 in BRCA1-proficient MEFs (Extended Data Fig. 3h). We found that overexpression of SHLD1 and to a lesser extent THAP1 significantly potentiated PARPi-induced genomic instability (Fig. 3h). These results are consistent with the notion that excessive expression of SHLD1 can overtly suppress HR, which explains why its expression must be fine-tuned by THAP1-mediated transcriptional regulation.

### THAP1 inhibits DNA end resection and RAD51 nucleofilament formation

The precise function of SHLD1 within the 53BP1 pathway is unknown. Previous studies have implicated SHLD3 as the anchor that attaches the entire Shieldin complex to RIF1, enabling its recruitment to 53BP1-decorated chromatin ^11, 20, 53^. We therefore tested whether THAP1 loss would impact the recruitment of 53BP1, RIF1 and SHLD3 to sites of DNA damage. We found that foci formation for 53BP1, RIF1, and SHLD3 was not impaired in *Thap1^-/-^Brca1*^Δ*11*^ cells but rather, was somewhat enhanced (Fig. 4a, b and Extended Data Fig. 4a, b). In BRCA1-deficient cells, unmitigated activation of the 53BP1-RIF1-Shieldin pathway impairs HR by blocking DSB end-resection and RAD51 nucleofilament formation ^14–21, 24^. Indeed, we found that deletion of THAP1 rescued both irradiation-induced RPA foci and RAD51 nucleofilament formation in *Brca1*^Δ*11*^ MEFs to wild-type levels (Fig. 4c, d). Thus, loss of THAP1 promotes end-resection and HR proficiency in BRCA1-deficient cells, which may explain why doubly deficient cells acquire resistance to PARPi and cisplatin.

**Fig. 4.**
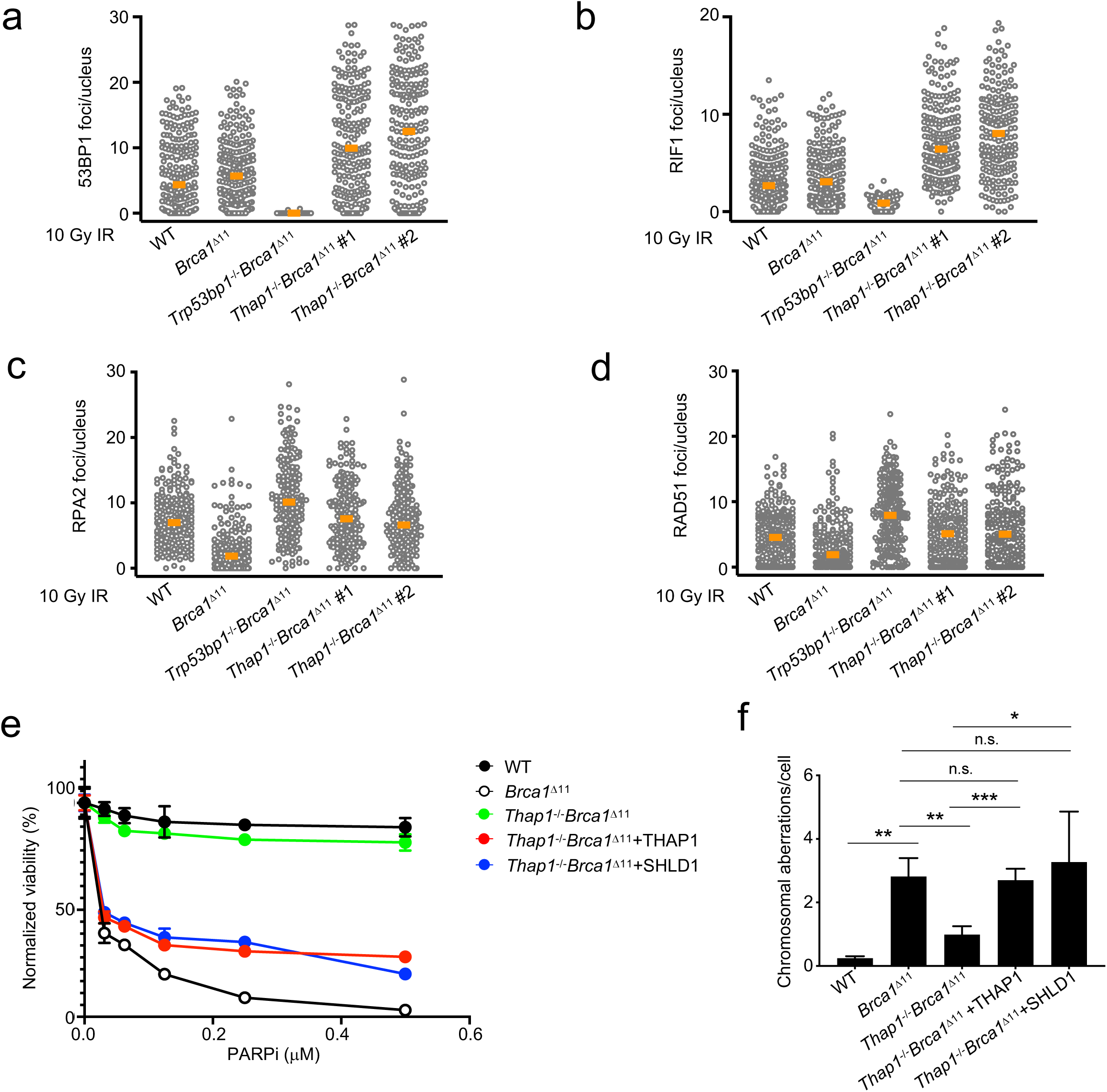
**THAP1-dependent *Shld1* expression inhibits HR and drives PARPi hypersensitivity in BRCA1-deficient cells.** (**a-b**) Quantification of 53BP1 (**a**) and RIF1 (**b**) foci in individual nuclei of *WT, Brca1*^Δ*11*^, *Trp53bp1^-/-^Brca1*^Δ*11*^ and two individual clones of *Thap1^-/-^Brca1*^Δ*11*^ MEFs. Cells were irradiated with 10 Gy and analyzed 1 h post-IR. (**c-d**) Quantification of RPA2 (**c**) and RAD51 (**d**) foci in individual EdU-positive (S-phase) nuclei of *WT, Brca1*^Δ*11*^, *Trp53bp1^-/-^Brca1*^Δ*11*^ and two individual clones of *Thap1^-/-^Brca1*^Δ*11*^ MEFs. Cells were irradiated with 10 Gy and analyzed 4 h post-IR. (**e**) Viability of *WT*, *Brca1*^Δ*11*^, *Thap1^-/-^Brca1*^Δ*11*^ and *Thap1^-/-^Brca1*^Δ*11*^ MEFs complemented with either THAP1 (*Thap1^-/-^Brca1*^Δ*11*^+THAP1) or SHLD1 (*Thap1^-/-^Brca1*^Δ*11*^+SHLD1) cDNA, as measured by CellTiter-Glo seven days after PARPi treatment. (**g**) Genomic instability (chromosome breaks and radials) detected in the metaphase spreads of *WT*, *Brca1*^Δ*11*^, *Thap1^-/-^Brca1*^Δ*11*^, *Thap1^-/-^Brca1*^Δ*11*^+THAP1, and *Thap1^-/-^ Brca1*^Δ*11*^+SHLD1 MEFs after 16 h of PARPi treatment (1 µM). At least 50 cells were scored per genotype and condition. The experiment was repeated five times. Statistical significance was determined by Welch’s t-test.

To provide definitive proof that the observed PARPi resistance in *Thap1^-/-^Brca1*^Δ*11*^ MEFs was due to decreased *Shld1* expression, we overexpressed SHLD1 in *Thap1^-/-^ Brca1*^Δ*11*^ MEFs (Extended Data Fig. 4c). Overexpression of SHLD1 in *Thap1^-/-^Brca1*^Δ*11*^ MEFs restored their hypersensitivity to PARPi (Fig. 4e). As expected, add-back of THAP1 in *Thap1^-/-^Brca1*^Δ*11*^ MEFs also restored PARPi hypersensitivity (Fig. 4e). Moreover, SHLD1 or THAP1 overexpression in *Thap1^-/-^Brca1*^Δ*11*^ MEFs increased chromosomal aberrations to levels comparable to or higher than in *Brca1*^Δ*11*^ parental cells (Fig. 4f). In contrast to SHLD1, ectopic SHLD3 expression did not promote PARPi hypersensitivity in *Thap1^-/-^Brca1*^Δ*11*^ cells, despite normal focal accumulation (Extended Data Fig. 4a, b, and d). Thus, THAP1-dependent *Shld1* transcription is essential for maintaining the functionality of Shieldin, but is not required for localizing the complex to DNA damage sites.

### THAP1 promotes Immunglobulin Class-Switch Recombination

Uncontrolled activation of the 53BP1-RIF1-Shieldin pathway compromises genome integrity during S/G2, but its function is uniquely required for NHEJ during class switch recombination (CSR), a G1-restricted genome rearrangement process that changes antibody effector functions ^15, 17, 18, 20, 54–56^. Notably, we found that loss of THAP1 severely compromised CSR in cytokine-stimulated CH12F3-2 mouse B cells, similar to inactivation of 53BP1, SHLD1 or SHLD3 (Fig. 5a). These results revealed a hitherto unknown physiological function THAP1 outside of the developing nervous system in which it maintains *Shld1* expression to promote NHEJ.

**Fig. 5.**
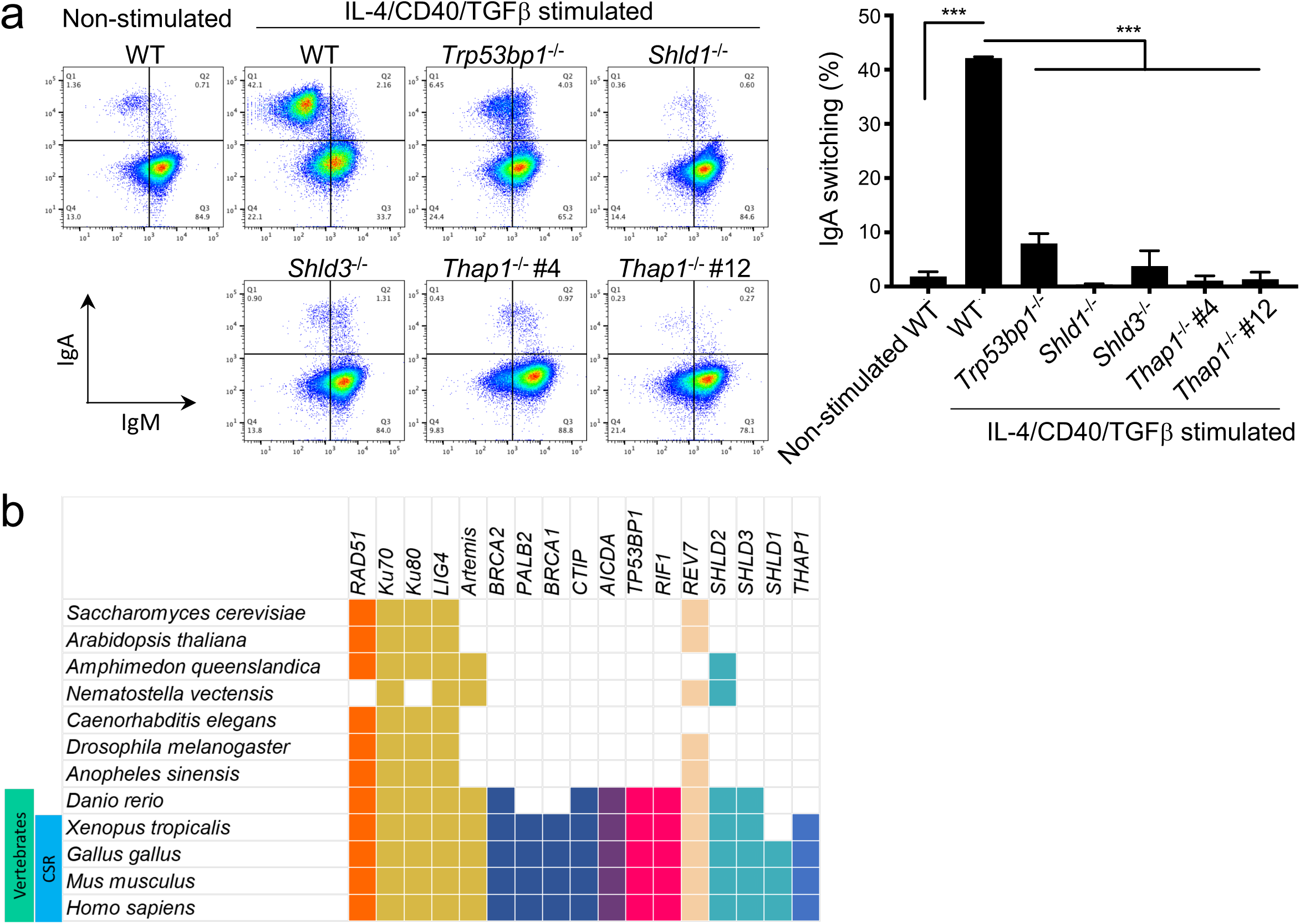
**THAP1 participates in physiological NHEJ.** (**a**) Representative flow cytometry plots of IgM-to-IgA class switch recombination (CSR) in *WT*, *Trp53bp1*^-/-^, *Shld1*^-/-^, *Shld3*^-/-^ and two individual clones of *Thap1*^-/-^ (#4 and #12) CH12-F3 cells 24 hours after cytokine stimulation (IL-4, CD40L and TGFb). Unstimulated WT cells are shown as a negative control. Quantification of IgM-to-IgA CSR is shown on the right and represents mean ± s.d., n=3. (**b**) Presence/absence table of selected genes in representative organisms, based on orthology data from the Ensembl database. Genes encoding THAP1, the Shieldin components and additional DSB repair factors are shown in relation to antibody CSR.

Although Shieldin genes are epistatic with 53BP1 and RIF1 with respect to CSR, they are phylogenetically younger than the upstream components of the DNA end-protection pathway ^18^. Indeed, it has been speculated that the evolution of Shieldin may have contributed to the emergence of a primitive form of CSR, with SHLD1 being the most recently evolved regulator ^18^. Interestingly, based on homology data across diverse eukaryotes, we found that THAP1 evolved recently with components of the Shieldin complex (Fig. 5b). It is therefore tempting to speculate that THAP1 might have co-evolved with Shieldin to regulate developmentally programmed NHEJ during CSR.

### THAP1 deficiency leads to unresolved DNA damage in the developing brain

Given previously established links between NHEJ deficiency and various neurodevelopmental abnormalities ^4^, we next investigated whether a perturbed DNA damage response could partially underlie the pathogenesis of DYT6. While germline deletion of murine *Thap1* is embryonic lethal ^44^, conditional deletion of *Thap1* in the central nervous system has been achieved using Nestin-Cre (*Thap1^flox/flox^Nestin-Cre*) ^49^). Re-analysis of RNA-seq dataset of *Thap1^flox/flox^ Nestin-Cre* mice showed that *Shld1* expression was significantly decreased in both the cerebellum and striatal tissues (Fig. 6a and Extended Data Fig. 5a), suggesting that loss of THAP1 may result in defective DNA repair in the brain.

**Fig. 6.**
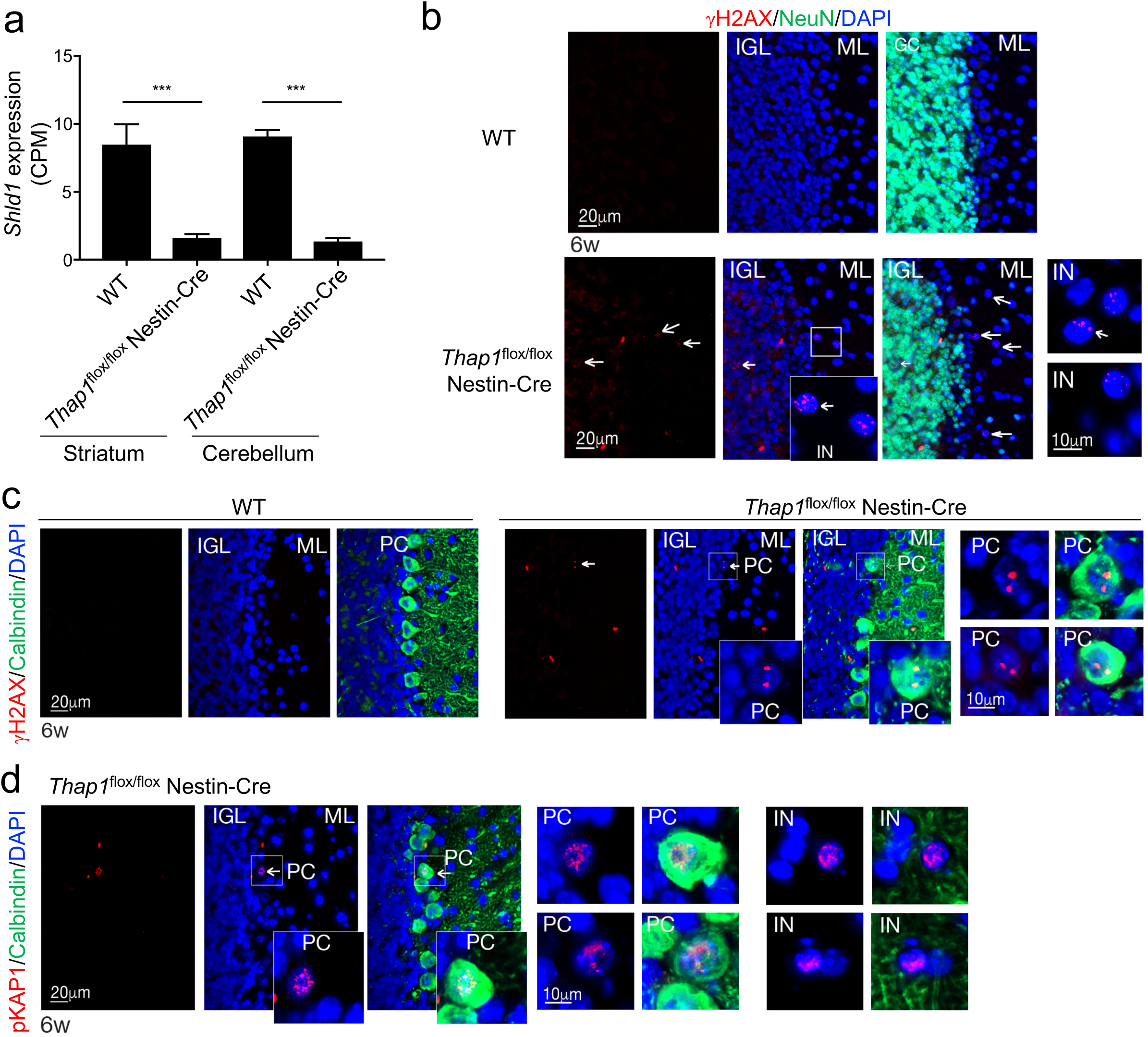
**THAP1-deficiency leads to ongoing DNA damage signaling in the brain.** (**a**) Analysis of *Shld1* gene expression in the striatum and cerebellum of *WT* and *Thap^flox/flox^Nestin-Cre* mice from a publicly available RNA-seq dataset (GSE123880). Data depict the mean value of counts per million reads (CPM) ± s.d.. (n = 3 for striatum and n = 4 for cerebellum, ***p<0.001). Statistical significance were determined by one-way ANOVA. (**b**) THAP1 loss causes genome instability in the cerebellum of 6-week-old *Thap1^Nes-cre^* mice. Unresolved DNA damage indicated by γH2AX (pSer139) is readily detectable in the inner granule layer (IGL) and in interneurons (IN) of the molecular layer (ML). The IGL is marked by NeuN immunopositivity. Expanded inset images show multiple nuclear puncta of γH2AX. (**c**) Purkinje cells (PC) marked by Calbindin immunopositivity show γH2AX nuclear puncta. (**d**) Phosphorylated KAP1 (pSer824)), a marker of DSB-induced ATM activation, is readily detectable in the nuclei of both PCs and interneurons (IN). Expanded inset images show multiple nuclear puncta of DNA damage-induced pKAP1 foci.

To explore this possibility, we generated an independently derived *Thap1^flox/flox^ Nestin-Cre* murine model, which exhibited motor abnormalities, including abnormal limb clasping during tail suspection similar to previous reports ^49, 57^. Histological examinations of various brain regions of 6-week old mice revealed significant increased immune-positivity of gH2ax and serine-824 phosphorylated Kap1 (p-Kap1) in THAP1-deficient cerebellum (Fig. 6b). For example, sporadic gH2ax immunoreactivity was observed throughout the inner granule layer of the of 6-weeks-old *Thap1^Nes-cre^* cerebellum, while in control tissue no DNA damage was found (Fig. 6b). Within the *Thap1^flox/flox^ Nestin-Cre* molecular layer, interneurons (Fig. 6b) also showed clear evidence of gH2ax as did calbindin positive Purkinje cells (PCs) (Fig. 6c). We also used p-Kap1, indicative of ATM activation, to confirm DNA damage accumulation (Fig. 6d). p-Kap1 foci showed a distribution similar to gH2ax with distinct, multiple individual nuclear foci present in *Thap1^flox/flox^ Nestin-Cre* neurons but not in controls, as shown for Calbindin positive PCs (Fig. 6d). Elsewhere in the *Thap1^flox/flox^ Nestin-Cre* brain, DNA damage accumulation was apparent in the cortex and hippocampal region as shown with gH2ax immunohistochemistry, while no sign of DNA damage was found in comparative regions of control tissue (Extended Data Fig. 5b, c). Thus, proper THAP1 function ensures neural genome stability in the brain.

## Discussion

The recently discovered Shieldin complex is a key enforcer of 53BP1-mediated DSB end protection, which blocks HR and confers chemotherapy hypersensitivity in BRCA1- deficient cells ^10, 11^. Yet, little is known about how Shieldin is functionally regulated *in vivo*. Here, we reveal that the evolutionarily conserved transcription factor THAP1 promotes transcription of the *Shld1* gene in mammalian cells. Our observation that loss of THAP1 restores HR and chemoresistance in both *Brca1*^Δ*11*^ (resection-deficient) and *Brca1*^Δ*11*^*Trp53bp1^S25A^* (RAD51 loading-deficient) MEFs is consistent with a critical function of Shieldin pre- and post-resection ^25, 35^. Moreover, the THAP1-SHLD1 axis supports productive CSR in murine B lymphocytes and maintains genome stability in the developing mouse brain. THAP1 acts in concert with co-regulators HCF1 and YY1, previously unknown to function in 53BP1-end protection pathways. Altogether, these observations highlight the need for cells to fine-tune Shieldin activity depending on the desired DSB repair outcome.

Shieldin complex components, particularly SHLD1 and SHLD3, are expressed at extremely low abundance compared to other DNA repair factors ^18, 31^. It is notable that a ubiquitious transcription factor module (THAP1-YY1-HCF1) would be specifically required to promote ultra low levels of *Shld1* expression. While additional studies are needed to determine whether the *Shld1* locus receives additional regulatory inputs to ensure its low steady-state transcriptional output, our data suggest that relatively low levels of 53BP1-RIF1-Shieldin activity may suffice for physiological NHEJ and that hyper-activation of this pathway could disproportionately inhibit HR in replicating cells. Consistent with this idea, we have demonstrated earlier that the efficiency of CSR was unaffected by increased 53BP1 expression or 53BP1 heterozygosity, whereas 53BP1 overexpression clearly exacerbated genomic instability induced by PARPi treatment not only in the BRCA1-deficient setting but also in BRCA1-proficient cells ^58^. Intriguingly, organisms with no clear orthologs for 53BP1, RIF1 or Shieldin also generally lack a recognizable orthologous *Brca1* gene (Fig. 5b). Given that excessive end protection can potentially compromise genome stability, we propose that orthologs of BRCA1 might have evolved specifically to counteract 53BP1-RIF1-Shieldin activity, thereby balancing the requirement for inherently error-prone NHEJ against the overarching need for maximizing genome stability through high-fidelity DSB repair.

Surprisingly, unlike BRCA1-deficient MEFs and human cells, in which loss of THAP1 largely rescued PARPi-induced genomic instability, we observed the opposite effect in BRCA2-mutant mESCs. Accordingly, clinical data suggested that low expression of THAP1 correlated with better outcome in *BRCA2*-mutated patients with serious ovarian cancer (Fig. 2d). Consistently, we found that low SHLD1 expression also correlates with improved survival in BRCA2 deficient ovarian cancer (Extended Data Fig. 2b). We note that a similar observation has previously been made in mouse cells lacking PALB2, in which the existing vulnerability to PARPi was further exacerbated upon loss of 53BP1 ^59^. Therefore, HR restoration in cells with an inhibited 53BP1-RIF1-Shieldin pathway seems to be contingent on RAD51 nucleofilament assembly. The role of BRCA1 in loading RAD51 can be partly compensated for by RNF168 ^60^, whereas PALB2 and BRCA2 are absolutely required ^25, 35^. Productive assembly of the recombinase has been shown to limit the extent of end resection ^61, 62^. Therefore, it is possible that deletion of THAP1/SHLD1 or 53BP1 would lead to excessive single-strand DNA in BRCA2-deficient cells, which in turn could increase genomic instability by engaging highly mutagenic RAD51-independent single strand annealing pathways ^52^.

Clinically, loss-of-function mutations in THAP1 cause the autosomal dominant dystonia disorder DYT6 ^32–34^. Employing a mouse model of DYT6, we demonstrated for the first time that THAP1 deficiency leads to unresolved DNA breaks in the developing mouse brain, particularly in the cerebellum. These DNA DSBs could arise from unrepaired oxidative DNA lesions that occur at preferentially at the promoter regions of neuronal genes during normal cellular aging ^63, 64^). NHEJ, which is promoted by Shieldin-mediated end protection, is likely to be the dominant DSB repair pathway in post-mitotic neurons. Therefore, it is plausible that impairment of THAP1-dependent *Shld1* expression could result in the accumulation of neuronal DSBs. In that context it is notable that ATM mutations, which abrogate 53BP1-Shieldin activity frequently present as generalized dystonia ^65, 66^. While additional studies are needed to test the role of SHLD1 in neurodevelopmental DNA repair, we speculate that unresolved DNA damage in the nervous system could be an important contributing factor to dystonia.

## Acknowledgements

We thank Sergio Ruiz Macias, Eros Lazzerini Denchi, and Nussenzweig lab members for stimulating discussions; Daniel Durocher for the *TP53^-/-^BRCA1^-/-^* human RPE1 cell line; Panagiotis A. Konstantinopoulos for assistance with TCGA analysis; K.S. was supported by a fellowship from the Uehara Memorial Foundation; Sequencing were supported from the NCI CCR Genomics Core; Flow cytometry analyses and sorting were performed at the NCI CCR LGI Flow Cytometry Core; The A.N. laboratory is supported by the Intramural Research Program of the NIH, an Ellison Medical Foundation Senior Scholar in Aging Award (AG-SS- 2633-11), the Department of Defense Idea Expansion (W81XWH-15-2-006) and Breakthrough (W81XWH-16-1-599) Awards, the Alex Lemonade Stand Foundation Award, and an NIH Intramural FLEX Award.

## Author Contributions

K.S., D.Z., E.C., and A.N. conceived and planned the study. K.S., D.Z., E.C., L.C.D., N.W., M.I., designed and performed experiments; K.S., D.Z., E.C., W.W. and F.B. analyzed the data; P.A.K. analyzed TCGA databases; M.E. and P.J.M. supervised and provided advice; A.N. supervised the study. K.S., D.Z., and A.N. wrote the manuscript with comments from the authors.

## Extended Data Figure Legends

**Extended Data Fig. 1.**
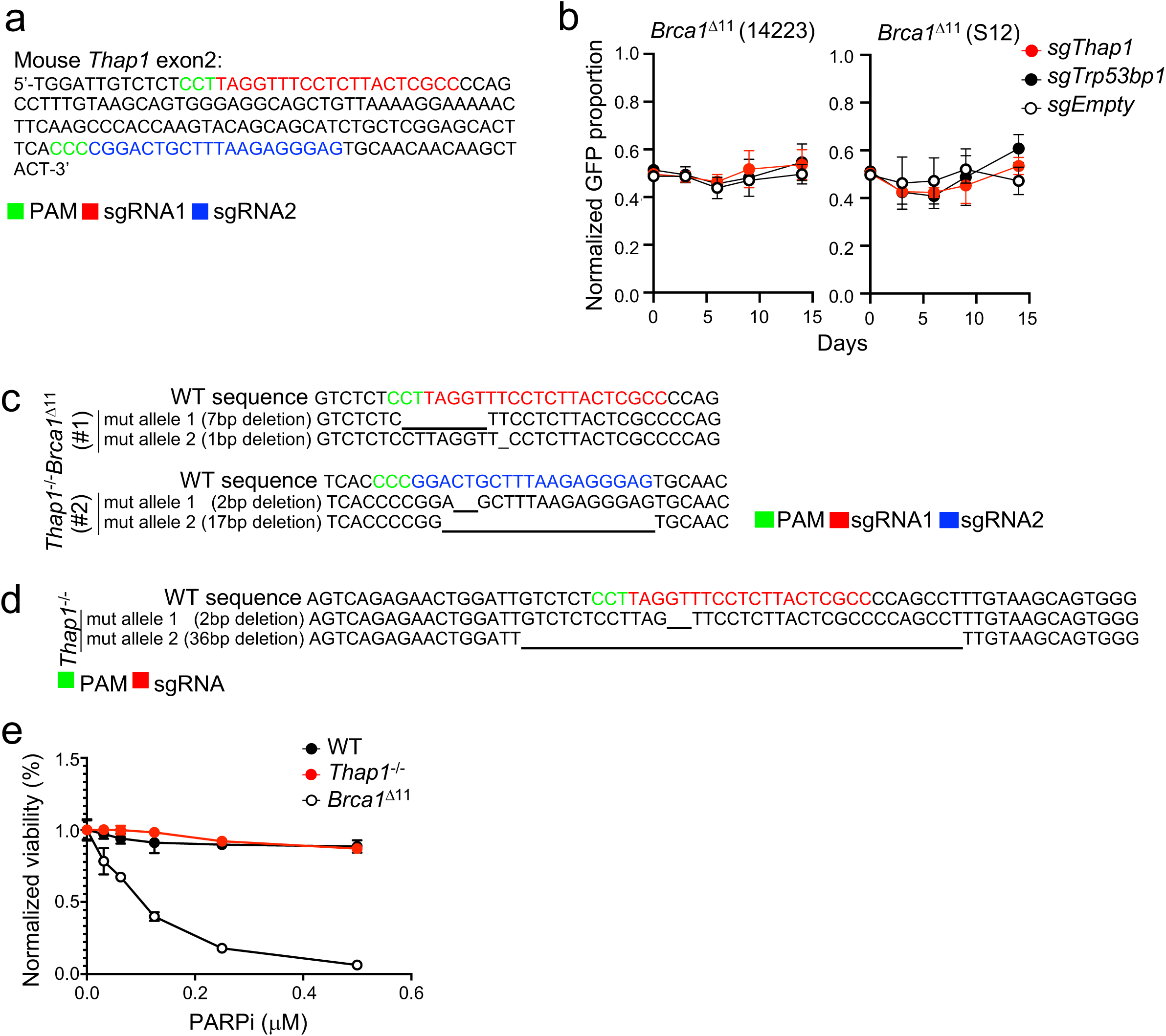
**Generation and characterization of *Thap1* knockout cells.** (**a**) Genomic sequences within exon 2 of the mouse *Thap1* gene. Two independent sgRNAs used to generate *Thap1* knockouts (sgRNA1;red and sgRNA2;blue) are shown along with their respective PAM sequences (green). (**b**) Multicolor Competition Assay (MCA): *Cas9^+^Brca1*^Δ*11*^ MEFs (14223 and S12) transduced with a specific guide RNA targeting *Thap1*, *Trp53bp1* or an empty vector (sgEmpty, all GFP-positive) were co-incubated (1:1 ratio) with *Cas9^+^Brca1*^Δ*11*^ MEFs transduced with non-targeting guides (sgLacZ, mCherry-positive). Data represent mean fraction of GFP-positive cells ± s.d., normalized to day 0 (n = 3). Vehicle treated control for Fig. 1c are shown. (**c**) Confirmation of successful editing of the *Thap1* locus in two independent *Thap1*^-/-^ *Brca1*^Δ*11*^ MEF clones (#1 and #2) by either sgRNA1 (red) or sgRNA2 (blue). (**d**) Confirmation of successful editing of the *Thap1* locus in a *Thap1*^-/-^ MEF clone by sgRNA1 (red). (**e**) Viability of *WT*, *Thap1*^-/-^ and *Brca1*^Δ*11*^ MEFs, as measured by CellTiter-Glo seven days after PARPi treatment.

**Extended Data Fig. 2.**
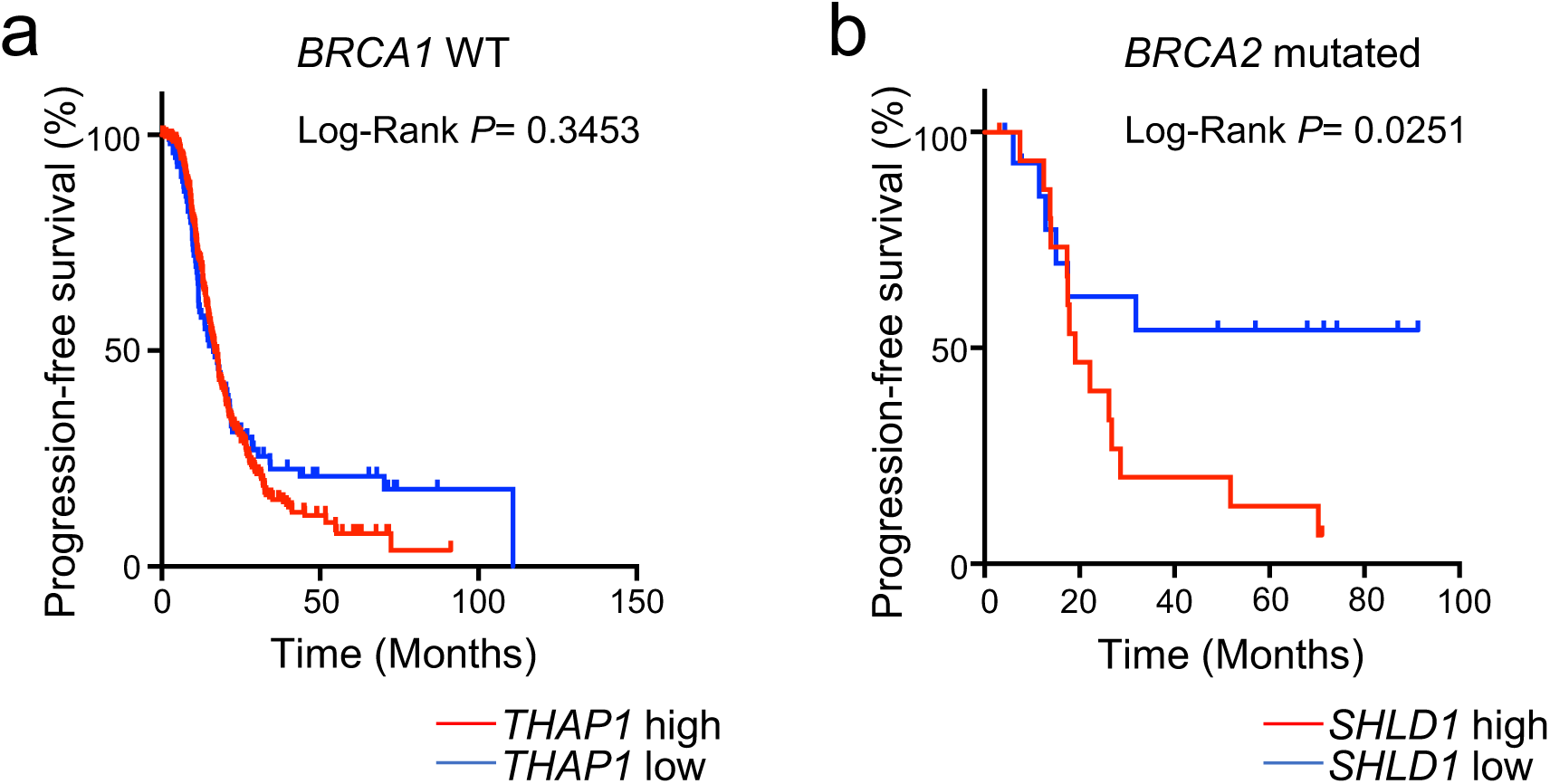
**Prognostic implications of THAP1 expression in ovarian serious adenocarcinoma.** (**a**) Progression-free survival (PFS) of *BRCA1*-WT ovarian serous adenocarcinoma patients on standard platinum-based regimens ^38^. Patients were defined as having *THAP1* low- or high-expressing tumors on the basis of the quintile of *THAP1* expression (z-scores < −0.67). PFS curves were generated by the Kaplan–Meier method. The difference between the PFS of *THAP1* low- versus *THAP1* high-expressing patients was assessed by the log-rank test (P > 0.05). (**b**) PFS of *BRCA2*-mutated ovarian serous adenocarcinoma patients on standard platinum-based regimens ^38^. The median expression of *SHLD1* for all tumors in this cohort was used to separate patients into *SHLD1* low- or high-expressors. The difference between the PFS of *SHLD1* low- versus *SHLD1* high-expressing patients was assessed by the log-rank test (P < 0.05).

**Extended Data Fig. 3.**
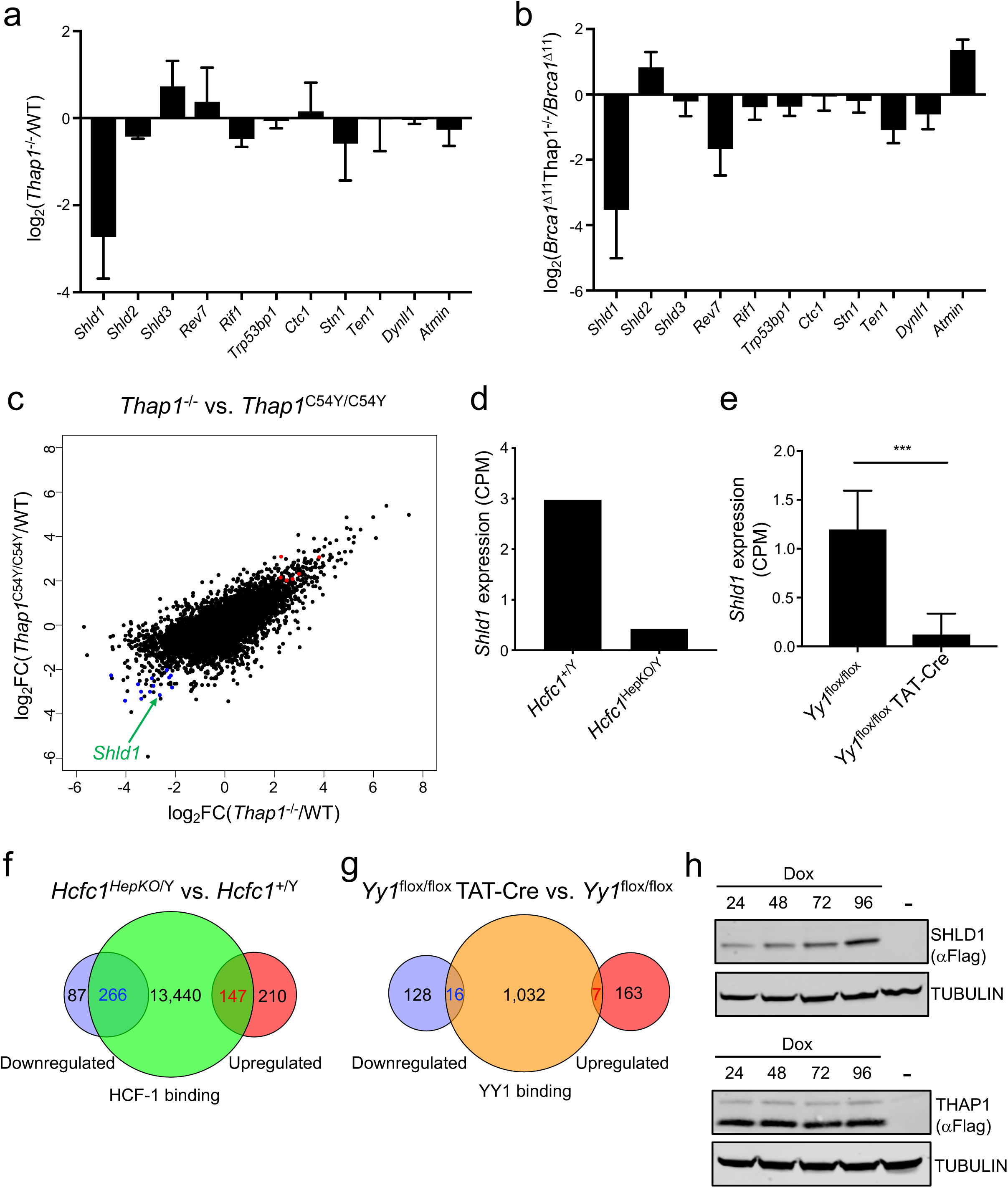
**Analysis of nascent RNA-seq, RNA-seq and ChIP-seq datasets for THAP1, HCF1, and YY1.** (**a** and **b**) Nascent RNA-seq showing relative differences (log2 fold-change) in the expression (CPM, counts per million mapped reads) of 53BP1 pathway-related genes in *Thap1^-/-^* versus WT MEFs (**a**) or *Thap1^-/-^Brca1*^Δ*11*^ versus *Brca1*^Δ*11*^ MEFs (**b**). Data represent mean relative difference ± s.d. (n = 3). (**c**) Scatter plot depicting relative differences (log2 fold-change) in gene expression (CPM) in *Thap1*^C54Y/C54Y^ versus *WT* and *Thap1*^-/-^ versus *WT* mESCs (RNA-seq, GSE86911) in relation to THAP1-bound genes (ChIP-seq, GSE86911). Genes that were bound by THAP1 and were either downregulated or upregulated in THAP1-deficient cells (both *Thap1*^C54Y/C54Y^ and *Thap1*^-/-^) are shown in blue and red, respectively. *Shld1* is indicated by the arrow. (**d**) Levels of *Shld1* gene expression in *Hcfc1^+/Y^* and *Hcfc1^HepKO/Y^* hepatocytes. Data are from a publicly accessible RNA-seq dataset (GSE115768) and represent mean CPM ± s.d. (n = 2). (**e**) Levels of *Shld1* gene expression in *Yy1^flox/flox^* and *Yy1^flox/flox^* TAT-Cre transfected B cells. Data are from a publicly accessible RNA-seq dataset (GSE145161) and represent mean CPM ± s.d. (n = 3, *p<0.05). Statistical significance was determined by the unpaired two-tailed Student’s t test. (**f**) Venn diagram depicting differentially expressed genes (log2 fold-change >2 and FDR <0.05) in *Hcfc1^+/Y^* versus *Hcfc1^HepKO/Y^* hepatocytes (RNA-seq, GSE115768) in relation to HCF1-bound genes in mESCs (ChIP-seq, GSE36030). The number of genes that were previously shown to be bound by HCF1 and were either downregulated or upregulated in HCF1-deficient cells are shown in blue and red, respectively. (**g**) Venn diagram depicting differentially expressed genes (log2 fold-change >2 and FDR <0.05) in *Yy1^flox/flox^* versus *Yy1^flox/flox^* TAT-Cre transfected B cells (RNA-seq, GSE145161) in relation to YY1-bound genes (ChIP-seq, GSE145161). The number of genes that were previously shown to be bound by YY1 and were either downregulated or upregulated in YY1-deficient cells are shown in blue and red, respectively. (**h**) Western blot analysis of doxycycline-dependent expression of exogenous SHLD1 and THAP1 proteins in *WT* MEFs 24 to 96 hours after induction with doxycycline (Dox) as detected by anti-Flag antibody.

**Extended Data Fig. 4.**
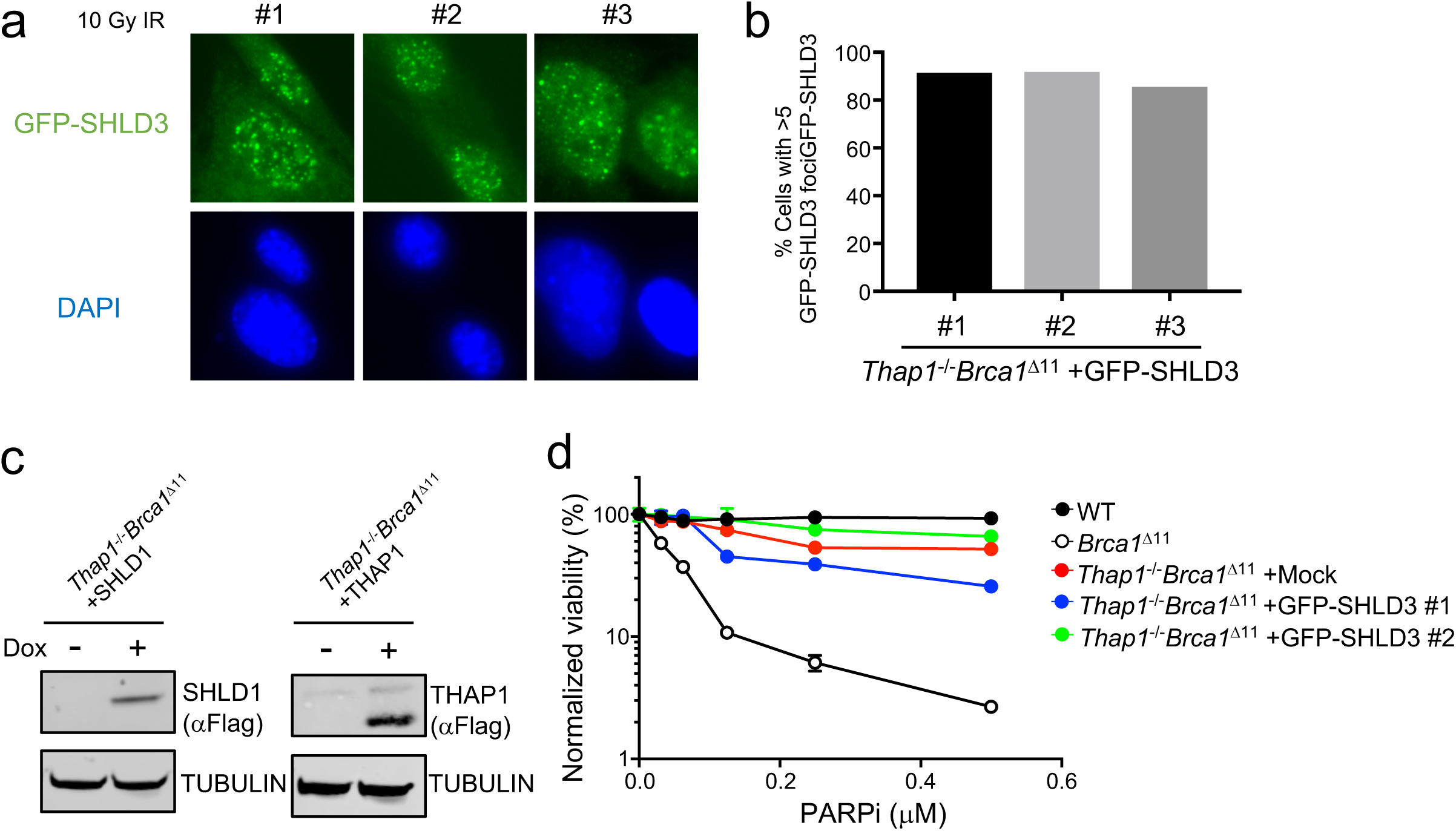
**Ectopic SHLD3 expression does not restore PARPi sensitivity in *Thap1*^-/-^*Brca1*^Δ11^ cells, despite normal focal accumulation.** (**a**) Representative immunofluorescence images depicting normal accrual of ectopically expressed GFP-SHLD3 in three independent *Thap1*^-/-^ *Brca1*^Δ*11*^ MEF clones. Cells were irradiated with 10 Gy and analyzed 1 h post-IR. (**b**) Quantification of the percentage of *Thap1*^-/-^ *Brca1*^Δ*11*^ MEFs containing >5 nuclear GFP-SHLD3 foci, as depicted in (**a**). (**c**) Western blot analysis of doxycycline-dependent expression of exogenous SHLD1 and THAP1 proteins in *Thap1*^-/-^ *Brca1*^Δ*11*^ MEFs 96 hours (SHLD1) and 72 hours (THAP1) after induction with doxycycline (Dox), as detected by anti-Flag antibody. (**d**) Viability of *WT*, *Brca1*^Δ*11*^, *Thap1*^-/-^*Brca1*^Δ*11*^ MEFs transduced with an empty vector (*Thap1*^-/-^*Brca1*^Δ*11*^+Mock) and two individual clones of *Thap1^-/-^Brca1*^Δ*11*^ MEFs complemented with SHLD3 cDNA (*Thap1^-/-^Brca1*^Δ*11*^+SHLD3#1 and *Thap1^-/-^ Brca1*^Δ*11*^+SHLD3#2), as measured by CellTiter-Glo seven days after PARPi treatment.

**Extended Data Fig. 5.**
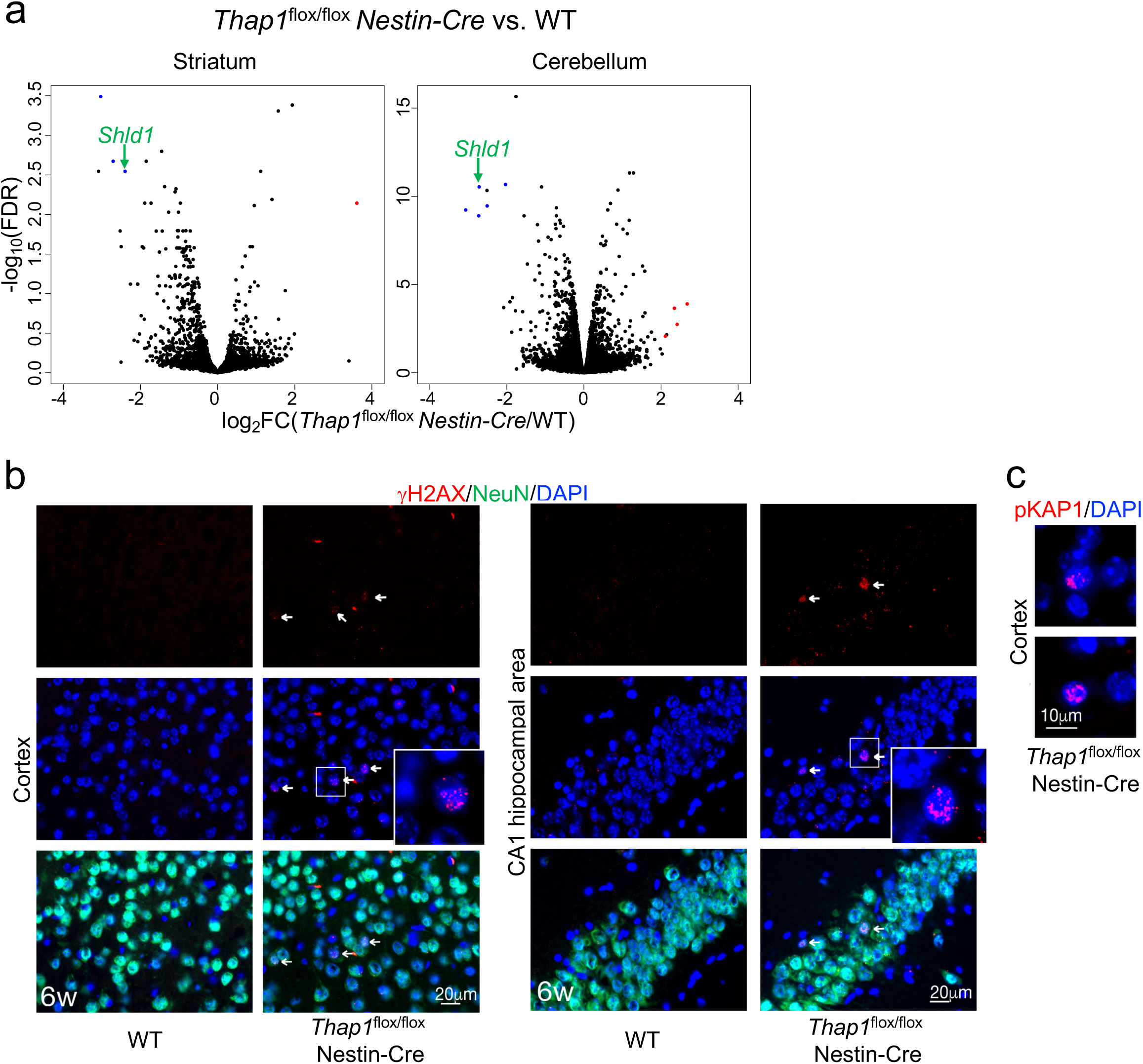
**Decreased *Shld1* expression and accumulation of unresolved DNA damage in the brain of *Thap1^flox/flox^ Nestin-Cre* mice.** (**a**) Volcano plot depicting relative differences (log2 fold-change) in gene expression (CPM) and the associated FDR in the striatum or cerebellum of *Thap1*^flox/flox^ *Nestin-Cre* mice as compare to WT. Data are from a publicly accessible RNA-seq dataset (GSE123880). Genes that were shown to be bound by THAP1 (ChIP-seq, GSE: GSE86911) and were either downregulated or upregulated in *Thap1^flox/flox^ Nestin-Cre* mice are shown in blue and red, respectively. (**b**) DNA damage resulting from THAP1 loss, as visualized by positive γH2AX immunostaining, is detected in multiple brain regions. In the 6-week-old *Thap1^Nes-cre^* cortex and CA1 area of the hippocampal region, γH2AX (pSer139) is observed as nuclear puncta sporadically in neurons as shown by NeuN immunostaining. (**c**) DNA damage, as indicated by phosphorylated KAP1 (pSer824), is also present sporadically throughout multiple brain regions including the cortex.

## Methods

### Cell lines and gene editing

Mouse embryonic fibroblasts (MEFs) were generated as previously described ^25^. The *Brca1*^Δ*11*^ and *Brca1*^Δ*11*^*Trp53bp1^S25A^* MEFs were generated from E13.5 embryos and grown in Dulbecco’s Modified Eagle’s Medium (DMEM, GIBCO) supplemented with 15% heat-inactivated fetal bovine serum (FBS, Gemini Bio-Products) and 1% penicillin + streptomycin (GIBCO). To establish immortalized MEF cell lines, primary MEFs between passages 2-4 were transiently transfected with a vector encoding SV40 T-antigen (pCMV-SV40T). SV40-immortalized MEFs were routinely cultured in DMEM supplemented with 10 or 15% FBS. For generation of Cas9-expressing *Brca1*^Δ*11*^ and *Brca1*^Δ*11*^*Trp53bp1^S25A^* MEFs, cells were transduced with the lentiCas9-Blast vector (Addgene #52962) and transductants were selected with blasticidin. Cells were then seeded at low densities (200–400 cells, depending on the cell line) on 15-cm dishes and single colonies were isolated using glass cylinders. Cas9 expression was confirmed by immunoblotting.

Generation of CRISPR knockout MEF and human cell lines were carried out as follows: Individual sgRNA (see Table S3 for sgRNA sequence) were cloned into LentiCRISPRv2 (Addgene #52961). To produce and package infectious lentivirus, HEK293T cells (American Type Culture Collection) at 80% confluency were co-transfected with LentiCRISPRv2-sgRNA, pRSV-Rev, pMDLg-pRRE and pHCMVG. After 72 h, viral supernatant was collected and passed through a 0.45 µm filter. Wild-type MEFs, *Brca1*^Δ*11*^ MEFs or *TP53*^-/-^*BRCA1*^-/-^ hTERT-RPE1 cells (1 × 10^5^) were seeded in 6- well plates and transduced using the filtered viral supernatant along with polybrene (10 μg/ml). RPE1-hTERT *TP53*^-/-^*BRCA1*^-/-^ and RPE1-hTERT *TP53*^-/-^ cells were kind gifts from Dr. Daniel Durocher (Lunenfeld-Tanenbaum Research Institute) and the packaging vectors were kind gifts from Dr. Shyam K. Sharan (NCI-Frederick, NIH). After 48 h, cells were subjected to puromycin selection (MEFs, 1.5 µg/ml; RPE1, 15 µg/ml) for five days. Thereafter, single cells were sorted into 96-well plates on a BD FACSAria UV instrument (BD Biosciences) and grown until colonies formed. Knockout clones were confirmed by Sanger sequencing of PCR amplicons of the targeted region that underwent editing. When different alleles of mutant clones had to be analysed, target regions were amplified by PCR using 50 ng genomic DNA in MyTaq Red mix (Bioline) according to manufacture instructions (primers are shown in Table S3). The amplicons were cloned using Zero Blunt TOPO PCR Cloning Kit (Thermo Fisher Scientific), and transformed into STBL3 chemically competent E. coli. (Thermo Fisher Scientific). Bacterial colonies containing plasmids with inserts were selected by colony-direct PCR, and were subsequently used for direct Sanger sequencing.

Generation of *Thap1*^-/-^*Brca2^Y3308X^* clones was performed as follows. Guide RNA against Thap1(see Table S3 for sgRNA sequence) was cloned into LentiCRISPRv2 with a Neo selection marker (Addgene #98292). *Brca2^Y3308X^* mES cells ^67^ were grown on 0.1% gelatin (Sigma) coated dishes in Dulbecco’s Modified Eagle’s Medium (DMEM, GIBCO) supplemented with 15% heat-inactivated fetal bovine serum (ESC-tested, Thermo Fisher Scientific), LIF (1000 U/ml) (Millipore), 0.1 mM MEM non-essential aminoacids, 1% glutamax and 55 µM b-mercaptoethanol. *Brca2^Y3308X^* mES cells (1 × 10^5^) cells were seeded in 6-well plates and transduced using virus produced from HEK293T cells along with polybrene (10 μg/ml). Thereafter, cells were selected with G418 (0.2 mg/ml). *Thap1*^-/-^ clones were confirmed by Sanger sequencing as described above.

*Trp53bp1*^-/-^, *Shld1*^-/-^, and *Shld3*^-/-^ CH12F3-2 cells were generated as previously described ^18^. *Thap1* mutant clones were edited through transient transfection with the pX330 (Addgene #42230) plasmid constructs expressing Cas9 and sgRNAs against *Thap1* (sgRNA: *Thap1*-sgRNA2, see Table S3), along with a plasmid encoding hCD4 (pcDNA3.1-Hygro-delta-hCD4), using an Amaxa Nucleofector II instrument (Lonza). Magnetic beads (STEMCELL Technologies 18052) were used to isolate hCD4^+^ cells 48 hour post-transfection. Thereafter, cells were grown for ∼7 days and subcloned by single cell sorting on a BD FACSAria UV instrument (BD Biosciences). *Thap1*^-/-^ clones were confirmed by Sanger sequencing as described above.

To generate GFP-SHLD3 expressing clones, *Thap1*^-/-^*Brca1*^Δ11^ MEFs were transduced with pCW-eGFP-SHLD3 (Addgene #114126) along with polybrene (10 μg/ml). Forty-eight hours after infection, the cells were treated with 1 µg/ml doxycycline for 24 h and single GFP^+^ cells were sorted into 96-well plates on a BD FACSAria UV instrument (BD Biosciences) and grown until colonies formed.

*WT* and *Thap1*^-/-^*Brca1*^Δ11^ MEFs expressing inducible exogenous THAP1 or SHLD1 were generated as follows. The PB-TRE-dCas9-VPR plasmid (Addgene #63800) was modified in which dCas9 was replaced by SnaBI, PacI and NotI restriction enzyme recognition sites. Thereafter, C-terminal Myc-Flag-tagged sgRNA target site-mutated version of the mouse *Thap1* cDNA (NM_199042) was inserted using SnaBI and PacI (PB-TRE-mTHAP1). Alternatively, C-terminal Flag-tagged mouse SHLD1 cDNA (NM_001358260.1) was inserted using SnaBI and NotI (PB-TRE-mSHLD1). sgRNA target site mutation in mouse *Thap1* cDNA was generated by Q5 Site-Directed Mutagenesis Kit (NEB, see Table S3 for primers). *WT* or *Thap1*^-/-^*Brca1*^Δ11^ MEFs (1 × 10^6^) were seeded in 10 cm dish and cultured for 24 hours. Cells were subsequently transfected with 7 µg of PB-TRE-mTHAP1 or PB-TRE-mSHLD1 along with 1.4 µg of a transposase expressing plasmid. Fourty-eight hours after transfection, cells were subjected to selection with 250 µg/ml hygromycin (Thermo Fisher Scientific) for 5 days. Cells were then seeded at low densities (200–400 cells) on 15-cm dishes and single colonies were isolated using glass cylinders. Selected clones were cultured in medium containing 1 µg/ml doxycycline for the indicated amount of time, and THAP1/SHLD1 expression was confirmed by immunoblotting.

### CRISPR–Cas9 screen

CRISPR–Cas9 screen was performed using genome-scale mouse Brie CRISPR knockout pooled library (Addgene #73633) ^68^. *Cas9^+^Brca1*^Δ*11*^ and *Cas9^+^Brca1*^Δ*11*^*Trp53bp1^S25A^* MEFs were transduced at a multiplicity of infection (MOI) of 0.3 and 400-fold coverage of the library. Cells were then selected with puromycin for 3 days before treatment with the clinically-approved inhibitor Olaparib (PARPi, 100nM) for a further 14 days. This dose corresponds with to the IC95 for the BRCA1-deficient cells. Surviving clones from each condition were collected, and genomic DNA (gDNA) was isolated (Blood & Cell Culture DNA Midi Kit, Qiagen) and subjected to PCR with Illumina-compatible primers, followed by Illumina sequencing. Genes enriched or depleted in the PARPi-treated samples were determined with the MAGeCK software package version 0.5.9.2.

### Multicolor growth competition assay (MCA)

One hundred thousand cells of two independent clones of Cas9-expressing *Brca1*^Δ*11*^ MEFs (14223 and S12) were infected with either virus particles of NLS-mCherry LacZ-sgRNA or NLS-GFP GOI-sgRNA (*Thap1*, *Trp53bp1* or the empty vector), respectively. Ninety-six hours after transduction, mCherry- and GFP-expressing cells were mixed 1:1 and 5 × 10^5^ cells were seeded with or without PARPi (Olaparib, 100 nM) in 100 mm dishes. During the course of the experiment, cells were subcultured when they approach confluency was reached. PARPi-containing medium was replaced every three days. Cells were analyzed on the day of initial plating (t = 0) and on days 3, 6, 9 and 14 using the flow cytometer (LSRFortessa; BD) and the percentage of GFP- and mCherry-positive cells were analyzed using FlowJo software (Tree Star). LIVE/DEAD discrimination was performed by staining with 1 µg/ml of 4’6-diamidino-2-phenylindole (DAPI, Invitrogen).

### Immunoblotting and Immunofluorescence

Western blotting was performed as described previously ^60^. Briefly, cells were collected and lysed in a buffer containing 50 mM Tris-HCl (pH 7.5), 200 mM NaCl, 5% Tween-20, 0.5% NP-40, 2 mM PMSF, 2.5 mM b-glycerophosphate (all from Sigma) and protease inhibitor cocktail tablet (complete Mini, Roche Diagnostics). Equal amounts of protein were loaded into precast mini-gels (Invitrogen) and resolved by SDS-PAGE. Proteins were blotted onto a nitrocellulose membrane, blocked with 5% membrane blocking agent (GE Healthcare) in TBS and incubated with the corresponding primary antibody. Primary antibodies were used at the following dilutions: anti-Flag (1:1000, Sigma) and anti-Tubulin (1:10,000, Sigma). Fluorescent secondary antibodies were used at a dilution of 1:15,000 (Li-Cor Biosciences). Detection of protein bands was performed by fluorescence imaging using a Li-Cor Odyssey CLx imaging system (Li-Cor Biosciences).

For immunofluorescence staining, MEFs were grown on 18 mm x 18 mm glass coverslips. Prior to g-irradiation (^137^Cs Mark 1 irradiator, JL Shepherd), cells were incubated with 10 µM EdU (Invitrogen) for 20 min. Following irradiation, cells were allowed to recover for 1 hour or 4 hours. Thereafter, cells were pre-extracted (20 mM HEPES, 50 mM NaCl, 3 mM MgCl2, 0.3 M sucrose, 0.2% Triton X-100) on ice for 5 min to remove soluble nuclear proteins. Extracted samples were fixed (4% para- formaldehyde), permeabilized (0.5% Triton X-100), incubated with the indicated primary antibodies followed by appropriate fluorochrome-conjugated secondary antibodies (Invitrogen). Next, click-IT chemistry was performed as per manufacturer’s instructions (Thermo Fisher Scientific) and DNA was counterstained with DAPI (Thermo Fisher Scientific). Images were captured at 63X magnification with an AxioCam MRc5 mounted on an Axio Observer Z1 epifluorescence microscope (Zeiss) or at 40x magnification on a Lionheart LX automated microscope (BioTek Instruments, Inc.). Quantification of nuclear foci and total nuclear intensity was performed using the Gen5 spot analysis software (BioTek). The antibodies used for standard immunofluorescence experiments were anti-RIF1 (1:5,000, gift of Davide Robbiani, Rockefeller University), anti-53BP1 (1:1000, Novus), anti-RAD51 (1:250, Abcam), anti-RPA (1:5,000, Abcam), and anti-GFP (1:500, Roche).

### Metaphase spreads and cell viability assays

MEFs and mESCs were treated with 0.5 µM or 1 µM PARPi (Olaparib, Selleckchem) for 16 hours, subsequently arrested at mitosis with 0.1 mg/ml colcemid (Roche) and metaphase chromosome spreads were prepared as previously described ^60^. Images were acquired using a Metafer automated scanning and imaging platform (MetaSystems). Clonogenic survival for a given treatment was calculated relative to the plating efficiency in non-treated controls. To determine cell growth and viability, MEFs were plated in 6- well plates (10,000 per well) and treated continuously with different doses of PARPi or cisplatin (Sigma) for 7 days. The drug-containing medium was replenished every three days and cells were subcultured when they approach confluency. On day 10, cell viability was determined using the CellTiter-Glo Luminescent Cell Viability Assay (Promega) as per manufacturer’s instructions.

### Nascent RNA-seq

Four million MEFs were labeled with 0.5 mM 5-ethynyl uridine (EU) for 30 min. Total RNA was extracted using TRIzol (Ambion). The NEBNext rRNA Depletion kit (human/mouse/rat) (New England Biosciences) was used to deplete rRNA from 1 µg of total RNA prior to sample biotinylation through Click-it reactions (Click-iT Nascent RNA Capture Kit, ThermoFisher C10365) as per the manufacturer’s specification. First-strand cDNA synthesis of the captured nascent RNA was done using the SuperScript VILO cDNA synthesis kit (Invitrogen), followed by AMPure XP purification (1.8X) and elution in (20 µl), Thereafter, second-strand cDNA synthesis was carried out in a total reaction volume of 30 µl that contained 0.6 mM dNTP, 1.2 mM of dUTP, 2 units of RNase H (Invitrogen) and 20 units of E. coli DNA polymerase I (Invitrogen), for 2.5 hr at 16°C. Double stranded cDNA was cleaned using 1.8X Agencourt AMPure XP beads and eluted in 20 µl of EB that was used for end-repair. End-repair was performed in 50 µl of T4 ligase reaction buffer containing 0.4 mM of dNTPs, 3 units of T4 DNA polymerase (NEB), 9 units of T4 Polynucleotide Kinase (NEB) and 1 unit of Klenow fragment (NEB) at 24°C for 30 min in a ThermoMixer C at 400 rpm. End-repair reaction was cleaned using 1.8X Agencourt AMPure XP beads and eluted in 15 µl of EB that was used for A-tailing reaction in 30 µl of NEBNext dA-Tailing reaction buffer (NEB) with 7.5 units of Klenow fragment exo- (NEB) at 37°C for 30 min. The 30 µl of the A-tailing reaction were mixed with Quick Ligase buffer 2X (NEB), 3,000 units of Quick ligase and 5 nM of annealed adaptor (Illumina truncated adaptor) in a volume of 50 µl and incubated at 25°C for 20 min. The adaptor was prepared by annealing the following HPLC-grade oligos: 5′-Phos/GATCGGAAGAGCACACGTCT-3′and 5′-ACACTCTTTCCCTACACGACGCTCTTCCGATC∗T-3′ (∗phosphorothioate bond). The ligation reaction was terminated by adding 50mM of EDTA and cleaned with 1.8X Agencourt AMPure XP beads and eluted in 15 µl of EB. Thereafter, samples were treated with 0.5 units of Uracil-DNA glycosylase (Thermofisher) for 15 min at 37°C and used for PCR amplification in a 50 µl reaction volume containing 1 µM of TruSeq barcoded primer p5, AATGATACGGCGACCACCGAGATCTACACNNNNNNNNACACTCTTTCCCTACA CGACGCTCTTCCGATC*T, TruSeq barcoded primer p7, CAAGCAGAAGACGGCATACGAGANNNNNNNNGTGACTGGAGTTCAGACGTGT GCTCTTCCGATC*T, (NNNNNNNN represents barcode and * a phosphothiorate bond), and 2X Kapa HiFi HotStart Ready mix (Kapa Biosci-ences). The temperature settings during the PCR amplification were 45 s at 98°C followed by 15 cycles of 15 s at 98°C, 30 s at 63°C, 30 s at 72°C and a final 5 min extension at 72°C. PCR reactions were cleaned with Agencourt AMPure XP beads (Beckman Coulter), run on a 2% agarose gel and a smear of 200-500bp was cut and gel purified using QIAquick Gel Extraction Kit (QIAGEN). Library concentration was determined with KAPA Library Quantification Kit for Illumina Platforms (Kapa Biosystems). Sequencing was performed on the Illumina Nextseq500 (75 bp single end reads).

### ChIP-Seq

ChIP-seq was performed as described previously ^69^ with a rabbit polyclonal antibody against THAP1 (Proteintech, 12584-1-AP). Twenty million MEFs were fixed in fresh media by adding 37% formaldehyde (F1635, Sigma) to attain a final concentration of 1% and incubated at 37°C for 10 min. Fixation was quenched by addition of 1 M glycine (Sigma) in PBS at a final concentration of 125 mM. Cells were washed twice with cold PBS and pellets were snap frozen in dry ice and stored at −80°C. Fixed cell pellets were thawed on ice and resuspended in 2 ml of cold RIPA buffer (10 mM TrisHCl pH 7.5, 1 mM EDTA, 0.1% SDS, 0.1% sodium deoxycholate, 1% Triton X-100, supplemented with proteinase inhibitor (Complete Mini EDTA free, Roche)). Sonication was performed using the Covaris S220 sonicator at duty cycle 20%, peak incident power 175, cycle/burst 200 for 30 cycles of 60 s sonication and 30 s of pause at 4°C. Chromatin was clarified by centrifugation at 21,000 g at 4°C for 10 min and precleared with 80 µl prewashed Dynabeads protein A (ThermoFisher) for 30 min at 4°C. 40 μl of prewashed Dynabeads protein A were incubated with 10 μg of antibody in 100 μl of PBS for 30 min at room temperature under continuous mixing, washed twice in PBS for 5 min and added to 1 ml of chromatin followed by overnight incubation at 4°C on a rotator. Beads were then collected in a magnetic separator (DynaMag-2 Invitrogen), washed twice with cold RIPA buffer, twice with RIPA buffer containing 0.3 M NaCl, twice with LiCl buffer (0.25 M LiCl, 0.5% Igepal-630, 0.5% sodium deoxycholate), once with TE (10 mM Tris pH 8.0, 1mM EDTA) plus 0.2% Triton X-100, and once with TE. Crosslinking was reversed by incubating the beads at 65°C for 4 hr in the presence of 0.3% SDS and 1 mg/ml of Proteinase K (Ambion). DNA was purified using Zymo ChIP DNA clean and concentrator kit (Zymo Research) and eluted in 20 μl. The entire ChIP DNA was used to prepare Illumina sequencing libraries. End-repair was performed in 75 μl of T4 ligase reaction buffer, 0.4 mM of dNTPs, 4 U of T4 DNA polymerase (NEB), 13.5 U of T4 Polynucleotide Kinase (NEB) and 1.5 U of Klenow fragment (NEB) at 24°C for 30 min in a ThermoMixer C at 400 rpm. End-repair reaction was cleaned using 2X Agencourt AMPure XP beads and eluted in 16.5 μl of EB that was used for A-tailing reaction in 20 μl of NEBNext dA-Tailing reaction buffer (NEB) with 7.5 U of Klenow fragment exo-(NEB) at 37°C for 30 min. The 20 μl of the A-tailing reaction was mixed with Quick Ligase buffer 2X (NEB), 3,000 U of Quick ligase and 5 nM of annealed adaptor (Illumina truncated adaptor) in a volume of 50 μl and incubated at 25°C for 20 min. Ligation was stopped by adding 50 mM of EDTA and cleaned with 1.8X Agencourt AM- Pure XP beads and eluted in 15 µl of EB that was used for PCR amplification in a 50 µl reaction with 1 μM primers TruSeq barcoded primer p5, TruSeq barcoded primer p7, and 2X Kapa HiFi HotStart Ready mix (Kapa Biosciences). The temperature settings during the PCR amplification were 45 s at 98°C followed by 14 cycles of 15 s at 98°C, 30 s at 63°C, 30 s at 72°C and a final 5 min extension at 72°C. PCR reactions were cleaned with Agencourt AMPure XP beads (Beckman Coulter), run on a 2% agarose gel and a smear of 200-500bp was cut and gel purified using QIAquick Gel Extraction Kit (QIAGEN). Library concentration was determined with KAPA Library Quantification Kit for Illumina Platforms (Kapa Biosystems). Sequencing was performed on the Illumina Nextseq500 (75bp single-end reads).

### Class switch recombination (CSR) assay in CH12-F3 cells

The mouse B lymphocyte cell line CH12-F3 cells was cultured and stimulated with 250 ng/ml CD40L, 10 ng/ml IL4 and 1 ng/ml TGFb (CIT) to induce immunoglobulin class switching from IgM to IgA. For FACS analysis of CSR efficiency, live CH12 cells were labeled with FITC-conjugated anti-IgM (ThermoFisher Scientific) and PE-conjugated anti-IgA (Southern Biotech) antibodies. All analyses were performed on FACS CantoII (BD Biosciences).

### TCGA analysis

Publicly available TCGA ovarian serous cystadenocarcinoma data on cBioportal ^70, 71^ were queried for *THAP1* and *SHLD1* expression data. Other clinical characteristics and information on *BRCA1* and *BRCA2* gene or expression alteration were also extracted for the 316 ovarian carcinoma tumors in the cohort ^38^. In total, 38 carcinomas with a *BRCA1* mutation and 34 carcinomas harboring a *BRCA2* mutation were isolated from the cohort and used for further analysis. All calculations were performed using and Graph Pad Prism v.7.0.

### Analysis of orthologous genes between species

Orthology presnce/absence data is based on Ensembl orthology data versions 98, 99 and 100 including human Ensembl genes used to mine Ensembl orthologs. Drosophila and Nematostella genes that were found to be orthologs of the human genes were used to further mine Ensembl Metazoa for additional animal orthologs, and human genes were further used to mine Pan-compara orthologs which includes taxa not present in the standard Ensembl dataset. A minimum of 15% identity of both query gene and target gene were required to avoid false positives.

### Histology and immunodetection

For cryosectioning, paraformaldehyde-perfused tissue was cryoprotected in buffered 25% sucrose (w/v) solution. Brains were embedded in Tissue-Tek OCT compound (Sakura), and sectioned sagittally at 9 µm using a CM3050S cryostat (Leica). Prior to immunodetection, sections were subjected to antigen retrieval according to the manufacturer’s directions (HistoVT One, Nacalai Tesque). Immunodetection was done with the following primary antibodies: Calbindin (1:500, Sigma, C9848), phospho-KAP-1 (S824) (1:500, Bethyl, A300-767A), NeuN (1:500, Chemicon, MAB377) and phospho- H2AX-Ser-139 (1:200; Cell Signaling, #2577). For fluorescence immunodetection, FITC- and Cy3-conjugated secondary antibodies (Jackson Immunologicals) were used and counterstained with DAPI (Vector Laboratories).

### Previously published RNA-seq and ChIP-seq datasets used in this study

RNA-seq and ChIP-seq datasets for *Thap1*^-/-^ mESC were obtained from GSE86911^45^. RNA-seq dataset for cerebellum and striatal tissues of *Thap1^flox/flox^ Nestin-Cre* mice were obtained from GSE123880 ^72^. RNA-seq and ChIP-seq datasets for *Yy1^flox/flox^* TAT-Cre primary B cells were obtained from GSE145161^51^. ChIP-seq dataset for YY1 in mESC were from GSE68195 ^73^. RNA-seq dataset for hepatocytes of *Hcfc1^HepKO/Y^* mice were obtained from GSE115768 ^50^. ChIP-seq dataset for HCF1 in mESC were from the Mouse ENCODE Project (GSE36030) ^74, 75^.

### Statistical analyses

Unless indicated, all data are presented as individual replicates. The total number of replicates, mean and error bars are explained in the figure legends. The statistical tests (Student’s, Welch’s, one-way ANOVA and log-rank) and resultant P values (represented by asterisks) are indicated in the figure legends and/or figure panels and were calculated using GraphPad Prism and R software (ns = p > 0.05; * = p < 0.05; ** = p < 0.01; *** = p < 0.001; **** = p < 0.0001).

